# A motif-vocabulary model of CAR T-cell intracellular domains identifies determinants of immunophenotype differentiation

**DOI:** 10.64898/2026.02.01.700582

**Authors:** Conor Kelly, Reem Bahr, Wenjun Zhu, Keyvan Keyvanfar, Praeep Dagur, Stefan Cordes

## Abstract

Chimeric antigen receptor (CAR) T-cell efficacy depends critically on the costimulatory domain, which shapes downstream signaling and the immunophenotype of manufactured products. Despite mechanistic evidence that immune receptors function as motif-based signaling scaffolds, CAR engineering has focused on a narrow set of costimulatory domains—principally CD28 and 4-1BB—leaving much of the available signaling design space unexplored. Here, we screened 1,243 naturally occurring intracellular domains as costimulatory modules in an anti-CD20 CAR backbone in primary human CD8^+^ T cells and quantified construct enrichment across memory-differentiation and PD-1-defined immunophenotypic compartments. Using Eukaryotic Linear Motif (ELM) annotations, we analyzed motif-phenotype associations via complementary statistical approaches: Mann–Whitney screening and negative binomial regression identified ELM features associated with differential construct representation, while Dirichlet-Multinomial modeling—which properly accounts for the compositional structure of FACS-partitioned data—revealed that individual ELMs do not significantly alter phenotype distributions. This discrepancy indicates that single motifs primarily affect proliferation or survival rather than differentiation fate. In contrast, construct-level analysis using a leave-one-out compositional test identified specific costimulatory domains with significant phenotype-shifting effects, demonstrating that particular *combinations* of ELMs—rather than individual motifs—determine immunophenotype. These results suggest that CAR T-cell differentiation state is governed by the integrated output of multiple signaling motifs and provide a combinatorial framework for rational costimulatory domain engineering.

## Introduction

Chimeric antigen receptor (CAR) T-cell therapy has transformed cancer immunotherapy for relapsed or refractory B-cell malignancies, achieving overall response rates of 70–90% in B-cell acute lymphoblastic leukemia and 70–80% in aggressive B-cell lymphomas. ^1–3^ However, 40–60% of initial responders ultimately relapse, many within the first year after infusion. ^4;5^ Insufficient T-cell persistence is repeatedly identified as a central limitation, particularly as CAR T-cell therapy is extended to solid tumors, autoimmune diseases, ^6^ and transplant tolerance applications. ^7–9^ Durable remissions are strongly associated with the immunopheno-type of the infusion product: patients whose leukapheresis material is enriched for CD27^+^CD45RO^−^CD8^+^ T cells, an early-memory population, exhibit superior CAR T-cell expansion, persistence, and long-term tumor control. ^4;5^ These findings establish T-cell differentiation state of the infusion product as a critical determinant of therapeutic durability and raise the question of how CAR design itself might be used to bias manufactured products toward persistence-associated immunophenotypes.

T cell activation requires two signals: antigen recognition through the T cell receptor (signal 1) and costimulation through receptors such as CD28 (signal 2). ^10;11^ Signal 1 alone induces anergy or apoptosis; costimulation is required for proliferation, survival, and differentiation into functional effector or memory cells. Chimeric antigen receptors collapse this two-signal requirement into a single synthetic receptor, rendering costimulation independent of ligand engagement and coupling it directly to antigen recognition: the CD3*ζ* chain provides signal 1, and an incorporated costimulatory domain provides signal 2. ^12;13^ Critically, the choice of costimulatory domain shapes the immunophenotype of the manufactured CAR T-cell product. CD28-based CARs drive rapid expansion and effector differentiation, whereas 4-1BB-based CARs promote central memory phenotypes, enhanced oxidative metabolism, and superior long-term persistence. ^14;15^ Because infusion product immunophenotype predicts clinical durability, costimulatory domain engineering offers a potential lever for biasing CAR T cells toward persistence-associated states during manufacture.

Costimulatory domains are intrinsically disordered scaffolds whose signaling capacity is encoded not by tertiary structure but by short linear motifs (SLiMs), classified as Eukaryotic Linear Motifs (ELMs). ^16;17^ These motifs recruit specific adaptor proteins, kinases, and phosphatases via direct recognition of specific amino acid sequences, and the particular combination of motifs within a domain determines downstream signaling outputs. ^18;19^ In this sense, ELMs constitute a molecular vocabulary from which evolution has assembled diverse signaling sentences, each specifying a distinct cellular response. The effects of any motif on cellular signaling depend on context: the combination of co-occurring motifs, their spacing, phosphorylation state, and the expression levels of cognate binding partners all modulate signaling strength and kinetics. The functional consequences of this signaling, including persistence, exhaustion resistance, and proliferative capacity, are what ultimately determine therapeutic efficacy; immunophenotype of the infusion product serves as an accessible but imperfect proxy for these outcomes that can be measured during manufacture. ^4;5^

Costimulatory receptors evaluated clinically fall predominantly into two major families with distinct signaling mechanisms. The immunoglobulin superfamily (IgSF), which includes CD28 and ICOS, contains short sequence motifs recognized by adaptor proteins with SH2 and SH3 domains, enabling rapid recruitment of PI3K, GRB2, and Lck. ^20;21^ The tumor necrosis factor receptor superfamily (TNFRSF), which includes 4-1BB, OX40, and CD27, instead recruits TRAF adaptor proteins through TRAF-binding motifs. ^22;23^ These families engage downstream signaling pathways that exhibit different kinetics: SH2/SH3-mediated pathways are rapidly inducible and largely binary, whereas TRAF-mediated pathways exhibit slower kinetics and more graded activation. ^24^ Consistent with this, 4-1BB-based CARs promote slower transition from memory to effector states and favor less differentiated, persistence-associated immunophenotypes relative to CD28-based CARs. ^25;26^ Beyond these two well-characterized families, immune cells employ additional motif classes that remain largely unexplored in CAR engineering.

Cytokine receptors exemplify this diversity: through successive gene duplications during vertebrate evolution, a single ancestral receptor expanded into families that recruit overlapping adaptor toolkits (JAKs, STATs) through distinct arrangements of phosphorylation-dependent docking sites, yet drive divergent transcriptional programs and cell fates. ^27–30^ This evolutionary precedent, functional diversification through combinatorial rearrangement of shared signaling motifs, suggests that the space of clinically favorable CAR T-cell phenotypes may extend far beyond the two dominant costimulatory domains currently in clinical use.

One such unexplored class is serine/threonine (Ser/Thr) phosphatase-docking motifs, which provide an orthogonal signaling vocabulary. ^31;32^ Motifs recruiting PP2A, calcineurin, 14-3-3, and WW domain proteins are present across immune receptor intracellular domains, where they enable graded signal modulation rather than the binary switching characteristic of SH2/SH3 pathways.

Beyond the dominant CD28 and 4-1BB domains, only a handful of alternatives (principally ICOS, OX40, and CD27) have seen significant clinical evaluation. ^15;26^ Recent pooled screens of CAR T cells have begun to sample broader CCD space, but these efforts have largely employed synthetic domains combining canonical immunoreceptor motifs (ITAMs and ITIMs) rather than the full diversity of ELM combinations found in naturally occurring signaling domains. ^33–36^ No study has, to our knowledge, systematically analyzed a large library of evolutionarily assembled CCDs to map how specific ELM combinations shape CAR T-cell immunophenotype during ex vivo manufacture.

Because ELM-mediated interactions depend on primary sequence rather than folded structure, costimulatory domains from diverse receptors can be inserted in modular fashion into CARs while retaining their signaling logic. We queried Gene Ontology and protein structure databases to compile a library of intracellular domains from transmembrane signaling receptors, both native and foreign to T cells, as candidate costimulatory domains. Here, we screened 1,243 of these candidate costimulatory domains (CCDs) in an anti-CD20 CAR. We transduced primary human CD8^+^ T cells with this pooled CAR library and co-cultured them with and without CD20^+^ malignant B cells under conditions designed to provoke exhaustion. We then sorted CAR T cells by memory differentiation state (naïve, central memory, effector memory) and PD-1 expression, and used targeted sequencing to quantify costimulatory domain representation within each sorted population. We annotated each CCD with the ELMs it contains and first analyzed whether individual ELMs influence T-cell subset distribution and PD-1 status. We then analyzed whether CCDs, representing evolutionarily assembled combinations of ELMs, shift phenotype distributions.

Distinguishing true phenotype effects from proliferation artifacts requires complementary statistical approaches. Mann–Whitney *U* tests on Pearson residuals provide sensitive screening for any ELM-phenotype associations, but can detect differences in total construct abundance across phenotypes that may reflect differential proliferation or survival rather than differentiation. A Dirichlet-Multinomial model directly captures the compositional structure of FACS-partitioned data (the constraint that counts across phenotypes sum to a fixed total for each construct), testing whether ELMs shift the *distribution* of cells across phenotypes independent of total abundance. To assess whether specific *combinations* of ELMs drive phenotype, we additionally performed construct-level analysis comparing each CCD’s phenotype distribution to a leave-one-out population mean. Our results nominate several costimulatory domains not previously evaluated in CAR T cells for further clinical investigation, and suggest that rational design of novel ELM combinations represents a promising future direction.

## Methods

### Candidate costimulatory domain library design

To compile a comprehensive set of candidate costimulatory domains (CCDs), we used Gene Ontology (GO) annotations to identify all human proteins that are integral components of the plasma membrane (GO:0098752, GO:0016021, GO:0005887) with biological processes related to T cell activation, differentiation, proliferation, anergy, costimulation, cytokine production, or TCR signaling (GO:0050868, GO:2000524, GO:0002725, GO:0042130, GO:0002669, GO:0050863, GO:0045580, GO:0042129, GO:0031295, GO:0050852). ^37^ We also included proteins annotated to broader immune-regulatory processes: regulation of apoptosis, chemokine production, innate immune response, interferon-*γ* production, leukocyte migration, regulation of immune response, and tolerance induction (GO:0043066, GO:0032722, GO:0045089, GO:0032729, GO:0002687, GO:0050776, GO:0002507). For proteins identified through these GO queries, we queried UniProt to determine membrane topology (defining extracellular, transmembrane, and cytoplasmic portions) and extracted the amino acid sequences of their cytoplasmic regions, yielding 1,660 distinct CCD sequences. ^38^ After codon optimization for human expression, we ordered as pooled oligonucleotides from Integrated DNA Technologies the 1,243 CCDs whose coding length was ≤350 bp.

The resulting library includes CCDs from antigen receptors, cytokine receptors, integrins, and canonical costimulatory molecules (CD28, 4-1BB, OX40, ICOS), spanning dozens of motif classes not previously studied in CAR T cells.

**Figure 1:**
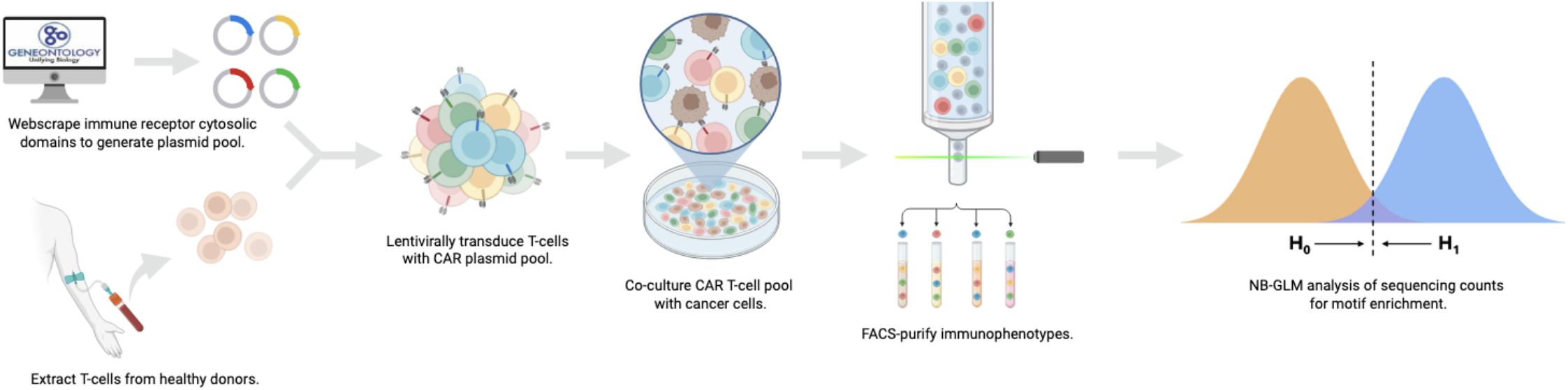
Graphical abstract. We queried GO and UniProt databases for candidate costimulatory domains and ligated codon-optimized oligonucleotides encoding these protein segments into an anti-CD20 CAR construct. Lentivirus was produced and used to transduce T cells from healthy donors. The resulting CAR T cells were co-cultured with CD20-expressing Raji cells for 10 days, then FACS-sorted into immunophenotypic subsets defined by memory differentiation state and PD-1 expression. Targeted sequencing quantified candidate costimulatory domain representation in each subset. Sequencing data were annotated by Eukaryotic Linear Motifs (ELMs) and analyzed using a negative binomial generalized linear model (NB-GLM).

### CAR T-cell production and screening

#### Synthesis of CAR plasmid pool

We used a third-generation anti-CD20 CAR comprising an anti-CD20 scFv, CD28 transmembrane domain, 4-1BB costimulatory domain, CD3*ζ* signaling chain, and truncated EGFR selection marker as the backbone for our constructs. ^39^ We excised the 4-1BB domain to linearize the plasmid. We converted the pooled single-stranded CCD oligonucleotides to double-stranded DNA by limited-cycle PCR and ligated them into the linearized backbone using NEBuilder HiFi DNA Assembly Master Mix (NEB, E2621), generating a pooled library in which each construct incorporated one CCD.

#### Lentiviral vector production

We produced lentiviral vectors by transfecting Lenti-X 293T cells (Takara Bio, 632180) with transfer plasmid and second-generation packaging plasmids psPAX2 (Addgene #12260) and pMD2.G (Addgene #12259) following established protocols. ^40^ We harvested viral supernatant at 24 and 48 h, concentrated it using Lenti-X Concentrator (Takara Bio, 631231), and stored aliquots at −80 ^°^ C.

#### T-cell isolation and transduction

We obtained venous blood from healthy adult donors under an IRB-approved protocol with informed consent. We enriched CD8^+^ T cells directly from whole blood using RosetteSep Human CD8^+^ T Cell Enrichment Cocktail (STEMCELL Technologies, 15023) and cryopreserved them. We thawed cryopreserved cells and activated them overnight with 500 IU/mL recombinant human IL-2 and Dynabeads Human T-Activator CD3/CD28 (Gibco, 11131D) prior to lentiviral transduction. We targeted a low multiplicity of infection (MOI) of 0.1–0.2 to ensure that the majority of transduced cells received a single CAR construct. On day 3, we removed beads by magnetic separation and expanded cells for 2 days with 500 IU/mL IL-2. On day 5, we stained cells with anti-EGFR-PE and enriched CAR^+^ cells using EasySep Release Human PE Positive Selection Kit (STEMCELL Technologies, 17654).

#### Co-culture conditions

We co-cultured enriched CAR T cells from two biological donors in triplicate (three technical replicates per donor) for 10 days under four conditions: (1) alone, (2) activated nonspecifically with anti-CD3/CD28 beads, (3) with irradiated K562 cells at a 1:1 effector:target ratio, or (4) with irradiated CD20^+^ Raji cells at a 1:1 effector:target ratio. We maintained cells in ImmunoCult-XF T Cell Expansion Medium (STEMCELL Technologies), refreshing medium and replenishing irradiated target cells every 2 days to maintain the 1:1 ratio.

#### FACS sorting

We stained CAR T cells and sorted them into six immunophenotypic populations defined by memory differentiation state and PD-1 expression. We first gated on lymphocytes by forward- and side-scatter, excluded doublets and dead cells, and then gated on CD3^+^CD8^+^ T cells. We identified CAR-expressing cells using an anti-IgG antibody (Jackson ImmunoResearch) that recognizes the IgG-derived hinge region of the CAR construct, and carried them forward for phenotypic analysis. We defined memory subsets by CCR7 and CD45RO expression: naïve (CD45RO^−^CCR7^+^), central memory (CM; CD45RO^+^CCR7^+^), and effector memory (EM; CD45RO^+^CCR7^−^). ^41^ Within each memory subset, we further stratified cells into PD-1^high^ and PD-1^low^ populations based on fluorescence intensity relative to isotype and fluorescence-minus-one controls, yielding six immunophenotypic fractions (Naïve_high_, Naïve_low_, CM_high_, CM_low_, EM_high_, EM_low_). ^42;43^ We performed six-way sorting on a BD FACSymphony S6 cell sorter, collecting live, singlet, CAR^+^ CD3^+^CD8^+^ lymphocytes and achieving ≥ 95% post-sort purity for all populations as assessed by re-analysis (Figure 2). We acquired additional activation and exhaustion markers (Supplementary Table 1) for post-sort characterization; these data will be reported separately.

**Figure 2:**
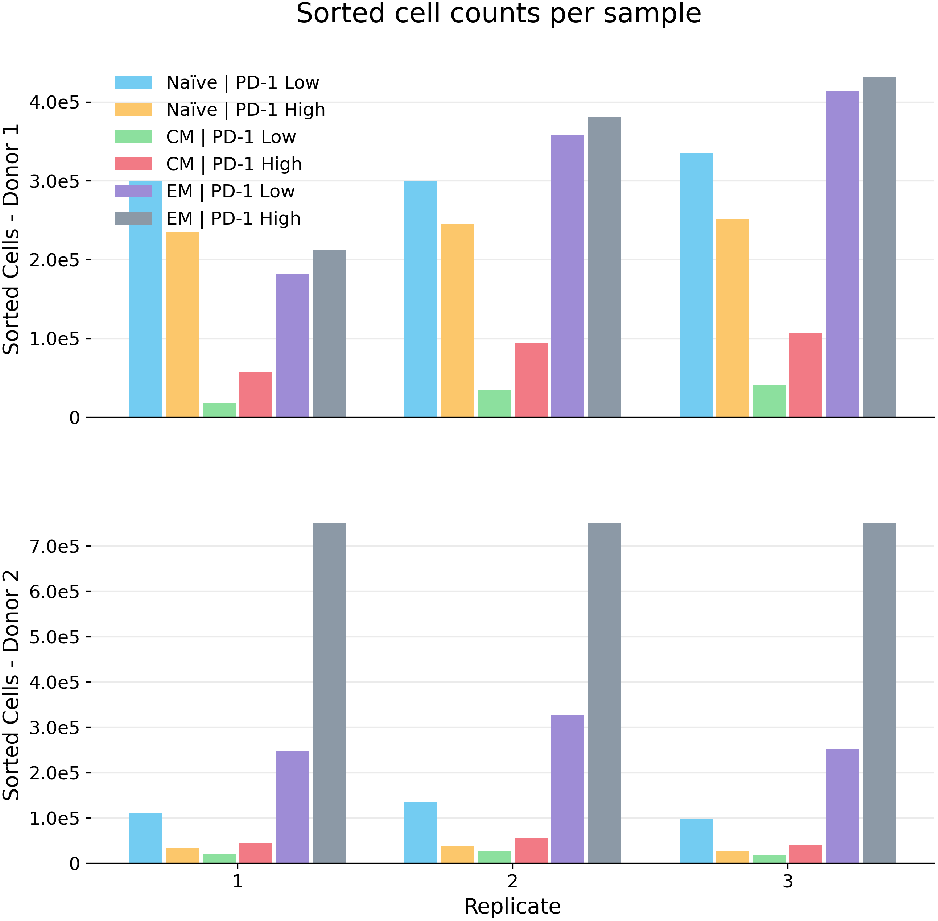
Sorted cell counts per sample across donors. Bar plots show the number of sorted CAR T cells recovered in each immunophenotypic fraction for each technical replicate from donor 1 (top) and donor 2 (bottom). Colors denote the immunophenotypic fractions (legend). The bottom x-axis labels indicate technical replicate number (1-3). Counts reflect post-sort recovery used for downstream sequencing and are provided to contextualize sampling depth across donors and replicates.

#### Targeted sequencing

We extracted genomic DNA from each sorted population using DirectPCR Lysis Reagent (Viagen Biotech, 301-C). We prepared sequencing libraries by two-step PCR: a first round amplified the candidate costimulatory domain with construct-specific primers, and a second round appended Illumina adapter sequences and sample indices. We demultiplexed FASTQ reads by sample index and mapped them to a reference of expected candidate sequences to produce a candidate-by-library count matrix (primary analysis pipeline: https://github.com/LIRGE/AnalyzeTargetedSequencing). Of 1,243 initial constructs, 1,071 yielded sufficient sequencing counts for downstream analysis. Secondary analysis code reproducing all figures is available as the costim screen package (Jupyter notebook included).

### Sequence annotation and statistical analysis

#### Terminology

We define the following terminology to describe experimental and analytical units. A *co-culture replicate* (CCR) is a single independent culture instance defined by the combination of donor, experimental condition, and technical replicate; each CCR represents an independent biological unit with shared transduction, expansion, and stimulation history. A *sequencing library* is the targeted PCR and sequencing product prepared from one FACS-sorted cellular fraction; we sorted each CCR into six phenotypically distinct T-cell subsets (Naïve, CM, and EM, each stratified by PD-1 expression), yielding six sequencing libraries per CCR. An *observation* is the raw integer read count *y*_*d,s*_ for candidate costimulatory domain *P* in sequencing library *Q*.

The full dataset comprised 144 sequencing libraries (2 donors

× 4 experimental conditions × 6 phenotypic fractions × 3 technical replicates). For inferential analyses, we focused on the antigenic condition (co-culture with CD20^+^ Raji cells), yielding 36 sequencing libraries from 6 CCRs (2 donors × 3 technical replicates). Each sequencing library contained read counts for 369 non-GPCR candidate costimulatory domains annotated with 49 normalized ELM groups. We encoded the ELM content of each candidate costimulatory domain *P* as a binary vector **x**_*d*_ ∈ {0, 1}^49^, where *R*_*d, j*_ = 1 indicates presence of ELM group *j* .

#### ELM annotation

We annotated each CCD with its constituent Eukaryotic Linear Motifs (ELMs). ^16^ ELMs are short, regular-expression-defined sequence patterns that represent the sequence prerequisites for interactions with specific adaptor proteins, kinases, and phosphatases. The presence of an ELM indicates the *potential* for interaction; whether the interaction actually occurs depends on additional factors including expression of cognate binding partners, post-translational modifications, and structural accessibility. This annotation creates a one-to-many mapping from CCDs to ELMs, enabling us to ask whether phenotype depends on individual motifs or on particular motif combinations.

To focus on signaling-relevant features, we restricted attention to docking (DOC) and ligand-binding (LIG) ELM classes, which mediate recruitment of adaptor proteins and enzymes. To analyze at the level of motif vocabularies and increase statistical power, we collapsed individual ELMs based on their shared interacting partner; for example, we merged multiple MAPK-binding docking motifs (e.g., <monospace>DOC_MAPK_DCC_ 7, DOC_MAPK_FxFP _2, DOC_MAPK_gen_1</monospace>) into a single functional group labeled “MAPK.” Filtering to groups present in at least 1% of domains yielded 49 normalized ELM groups. This collapsing was implemented to reduce dimensionality for NB-GLM fitting and assumes approximate functional equivalence among motifs recruiting the same partner; this assumption may not hold if different binding modes produce opposing effects, and analysis at the level of individual ELMs is ongoing.

#### Classification of CCDs by GPCR architecture

Not all intracellular domains from transmembrane proteins contain functional ELMs; motif-based signaling requires intrinsically disordered regions that expose linear motifs to cytoplasmic binding partners. G-protein coupled receptors (GPCRs) constitute a large fraction of our CCD library, but GPCR intracellular architecture differs fundamentally from single-pass receptors. The canonical seven-transmembrane GPCR contains four intracellular segments: intracellular loops 1 and 2 (ICL1, ICL2) are relatively short and conserved, forming structured interfaces that determine G-protein coupling specificity, ^44;45^ whereas intracellular loop 3 and the C-terminal tail (ICL3–CTail) are intrinsically disordered and enriched for signaling motifs, including phosphorylation sites that recruit arrestins and other modular signaling proteins. ^46;47^ Because our library treats each intracellular segment as a separate CCD, co-operative effects between segments (particularly ICL2’s role in modulating G-protein selectivity through conformational coupling with ICL3) ^48^ are not captured. We therefore analyzed GPCR-derived CCDs separately, with the expectation that ICL2 would show minimal motif-based signal (serving as an internal negative control) while ICL3–CTail segments would show intermediate signal.

We classified CCDs as GPCR or non-GPCR based on whether their source protein’s UniProt entry encodes a seven-transmembrane receptor. Of 1,071 retrieved constructs, 369 were non-GPCR and 702 were GPCR-derived (Figure 3).

**Figure 3:**
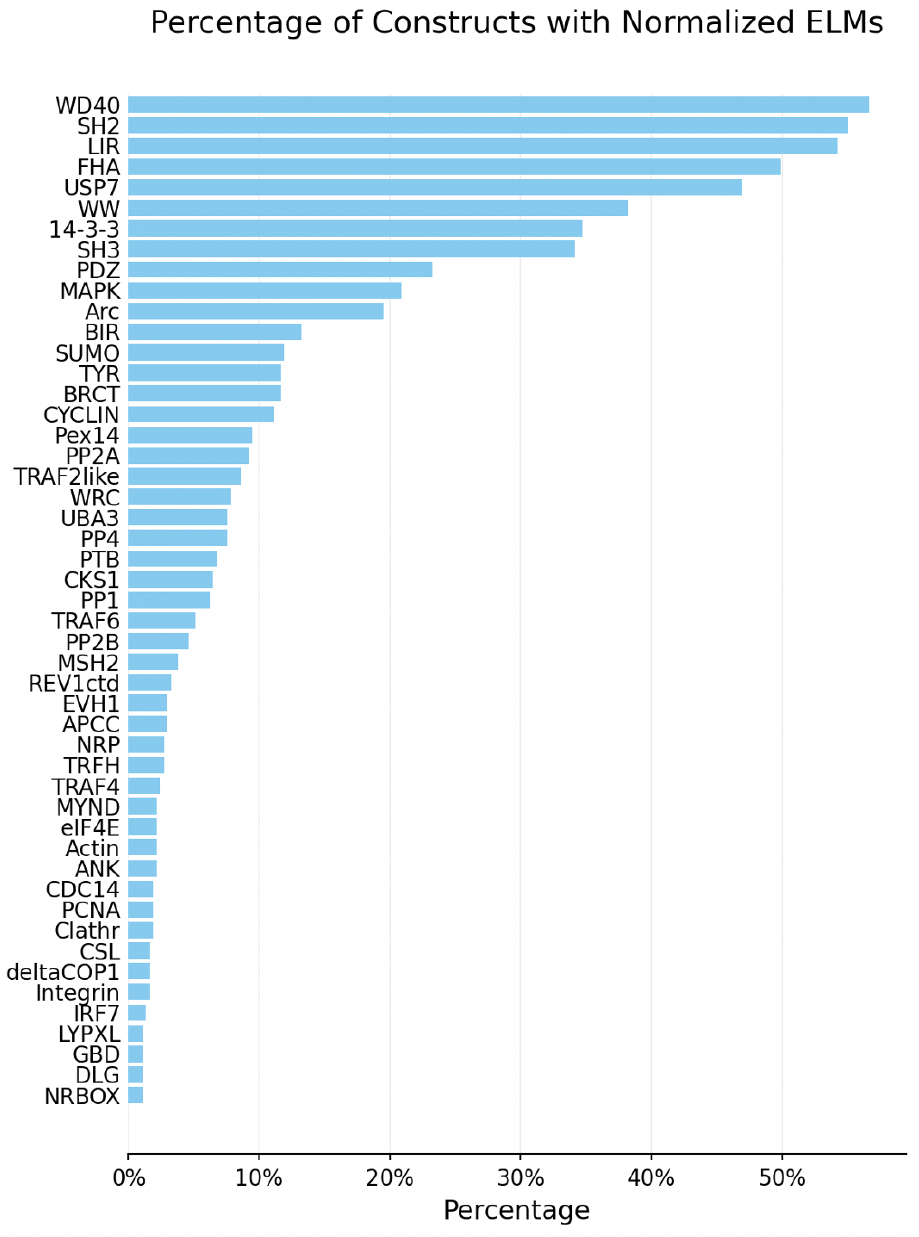
Normalized ELM coverage of the costimulatory-domain pool. Distribution of the 49 normalized ELM groups (DOC/LIG classes, collapsed by shared interacting partner and filtered to features present in ≥ 1% of domains) across the 369 non-GPCR costimulatory domains.

We analyzed non-GPCR and GPCR-derived CCDs separately to respect the potential context-dependence described above: when GPCR intracellular segments are isolated as individual CCDs, they lack the cooperative interactions that enable their native function.

For GPCR-derived CCDs, we further subdivided by intracellular segment: ICL2 versus ICL3 and the C-terminal tail (the fourth intracellular segment). ICL2, when isolated, lacks the conformational context needed for G-protein engagement and serves as an internal negative control. ICL3 and the C-terminal tail are more disordered and retain phosphorylation-dependent motifs that can function independently; we expected intermediate signal from this subset.

We applied Mann–Whitney screening and Dirichlet-Multinomial modeling to each structural category independently.

#### Mann–Whitney screening

For initial screening, we computed Pearson residuals from raw counts to remove library size effects:

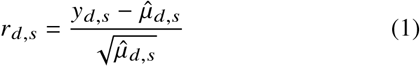

where 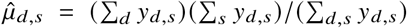 is the expected count under independence. Within each phenotype *p*, we tested whether candidate costimulatory domains containing ELM group *j* exhibited different residual distributions than those lacking the ELM using the two-sided Mann–Whitney *U* test:

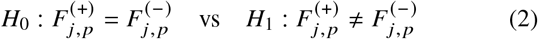

where 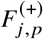 and 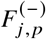 denote the residual distributions for domains with and without ELM *j*, respectively. We quantified effect sizes using Cliff’s delta (*δ*), a non-parametric measure bounded by [− 1, 1] representing the probability that a randomly selected domain with the ELM has a higher residual than one without, minus the reverse probability. To obtain pooled T-subset contrasts (e.g., EM vs CM averaged over PD-1), we combined phenotype-specific *p*-values using Fisher’s method and averaged Cliff’s delta values.

We adjusted *P*-values for multiple testing using the Benjamini–Hochberg procedure within each comparison and assessed significance at FDR *q* < 0.10 for screening.

#### Dirichlet-Multinomial analysis

For ELMs identified in Stage 1, we fit a Dirichlet-Multinomial (DM) model that directly captures the compositional structure of FACS data (the constraint that cell counts across the six phenotypic fractions sum to a fixed total for each CCD within each CCR). The model specification is:

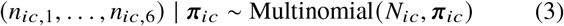

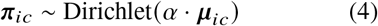

where *i* indexes CCDs, *c* indexes CCRs, *N*_*ic*_ = ∑_*p*_ *n*_*ic, p*_ is the total count, and *α* > 0 is the concentration parameter controlling overdispersion. The mean phenotype probabilities ***μ***_*ic*_ are modeled via a multinomial logit:

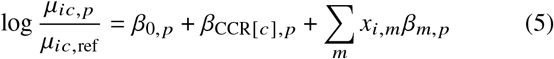

where *β* _*m, p*_ is the effect of ELM *m* on phenotype *p* relative to the reference (Naïve_Low_).

For inference, we computed phenotype contrasts using the full covariance matrix from the fitted model.

#### CCD-level compositional analysis

To test whether specific costimulatory domain constructs, representing particular combinations of ELMs, shift phenotype distributions, we performed a construct-level G-test (log-likelihood ratio test) comparing each CCD’s phenotype distribution to a leave-one-out population mean. For each CCD *i*, we computed:

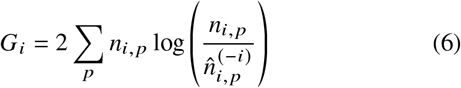

where *n*_*i, p*_ is the observed count in phenotype *p* and 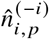 is the expected count based on the phenotype distribution of all other CCDs (leave-one-out mean). Under the null hypothesis of no phenotype shift, 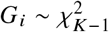 where *k* is the number of phenotypes.

We computed effect sizes for specific phenotype contrasts (EM vs CM, Naïve vs CM, Naïve vs EM, PD-1^low^ vs PD-1^high^) as log-odds ratios relative to the leave-one-out population. We adjusted *p*-values using Benjamini–Hochberg FDR correction.

#### Multiple testing correction

We adjusted *P*-values from all analyses for multiple testing using the Benjamini–Hochberg procedure within each phenotype comparison. For ELM-level MW screening, we assessed significance at FDR *q* < 0.10; for CCD-level analysis, we report results at FDR *q* < 0.10.

## Results

### Mann–Whitney screening identifies ELM-associated abundance differences

To screen for ELM features associated with differential construct representation, we applied Mann–Whitney *U* tests to Pearson residuals within each phenotype, comparing CCDs containing each ELM group to those lacking it. Effect sizes were quantified using Cliff’s delta; *p*-values were FDR-corrected within each comparison. For memory-state comparisons, we report pooled contrasts averaging effects across PD-1 levels. Throughout, contrasts are oriented as *A* − *B* (e.g., EM−CM): positive Cliff’s delta indicates higher abundance in phenotype *A*.

At FDR *q* < 0.10, Mann–Whitney screening identified 19 significant ELM-phenotype associations out of 147 tests across pooled T-subset comparisons (Figure 4).

**Figure 4:**
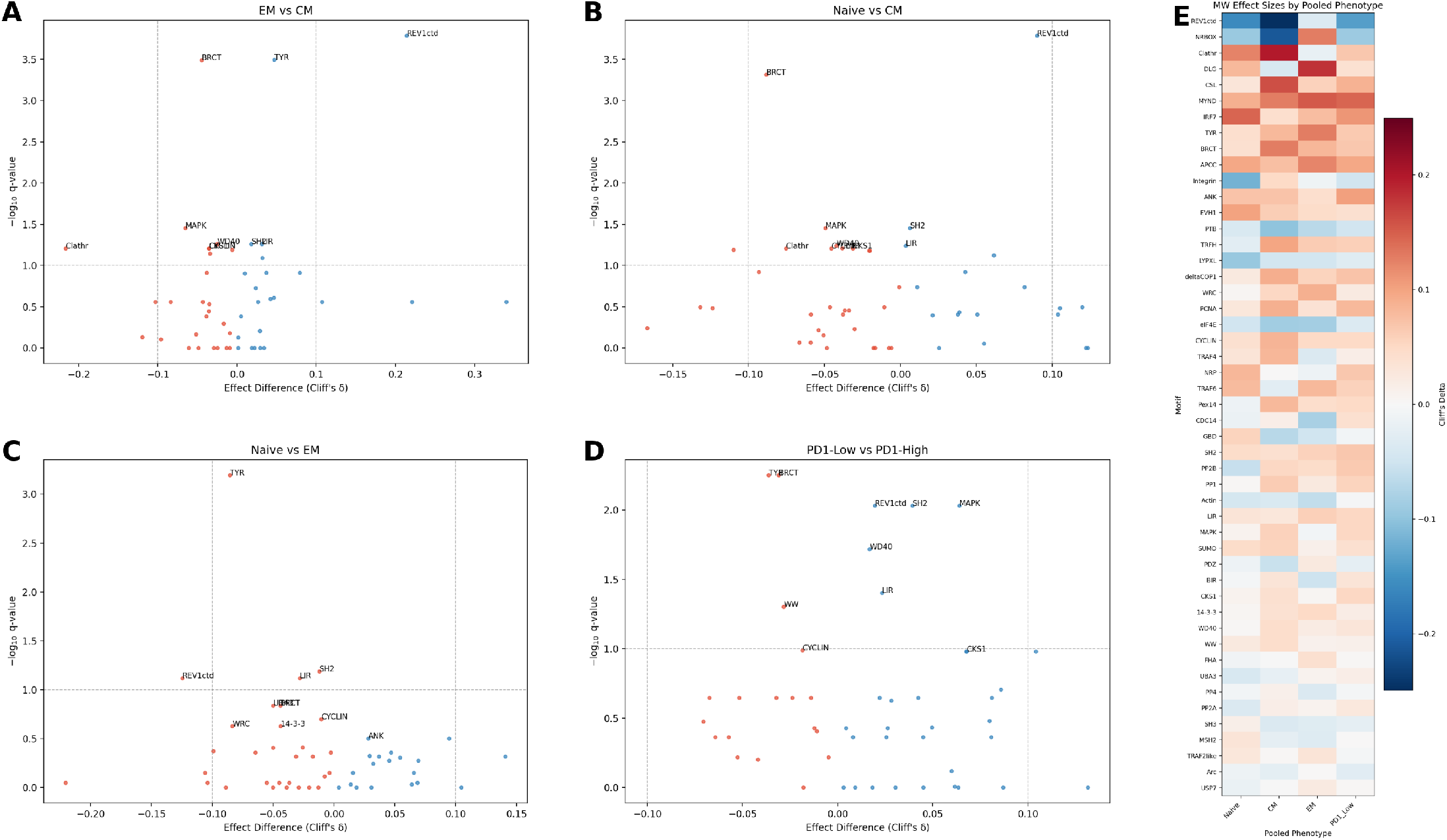
Mann–Whitney screening identifies ELM-phenotype associations in non-GPCR CCDs. Analysis restricted to ELMs within non-GPCR costimulatory domains, which include canonical immune receptors (4-1BB, CD28, OX40, etc.). (**A–D**) Volcano plots for pooled contrasts comparing ELM effects across immunophenotypes using Mann–Whitney *U* tests. Each point represents one normalized ELM group (*n* = 49). The x-axis shows Cliff’s delta (*δ*) for the contrast *A-B* (positive values indicate enrichment in phenotype *A*), and the y-axis shows statistical support as → log_10_(*q*), where *q* is the Benjamini-Hochberg FDR-adjusted *p* -value. Points colored by significance at FDR < 0.10. Contrasts shown: (A) EM vs CM, (B) Naïve vs CM, (C) Naïve vs EM, (D) PD-1^low^ vs PD-1^high^. (**E**) Heatmap of pooled Cliff’s delta values across phenotypes. Columns show T-subset effects pooled over PD-1 levels and PD-1 effects pooled over T-subsets. See Supplementary Figure 1 for parallel analysis of GPCR ICL3/C-tail domains, and Supplementary Figure 2 for the GPCR ICL2 negative control.

### Compositional analysis reveals no ELM-level phenotype effects

To test whether individual ELMs alter the *distribution* of cells across phenotypes, as opposed to affecting total construct abundance, we fit a Dirichlet-Multinomial (DM) model that directly captures the compositional constraint of FACS-partitioned data.

Strikingly, no ELM-phenotype associations reached significance at FDR < 0.10 in the DM framework (Supplementary Figure 3). This null result is informative: the discrepancy between MW screening (which identified many associations) and DM analysis (which identified none) indicates that individual ELMs affect construct *abundance* within phenotypic compartments (likely reflecting differential proliferation or survival) but do not alter the phenotype *composition* itself. In other words, the presence of a particular ELM may influence how many CAR T cells survive or expand, but does not determine whether those cells differentiate into central memory, effector memory, or naïve phenotypes.

### CCD-level analysis reveals combinatorial phenotype effects

If individual ELMs do not shift phenotype distributions, what does? To address this, we performed construct-level (CCD) analysis, testing whether specific costimulatory domains, each representing a particular combination of ELMs, shift phenotype distributions relative to the population.

Using a leave-one-out G-test, we identified numerous CCDs with significant phenotype-shifting effects at FDR < 0.10 (Figure 5). These included canonical costimulatory domains such as 4-1BB (TNFRSF9), as well as domains from interleukin receptors and other immune signaling proteins. The contrast between the ELM-level null result and the CCD-level positive findings indicates that phenotype specification arises from the integrated output of multiple signaling motifs rather than from any single ELM.

**Figure 5:**
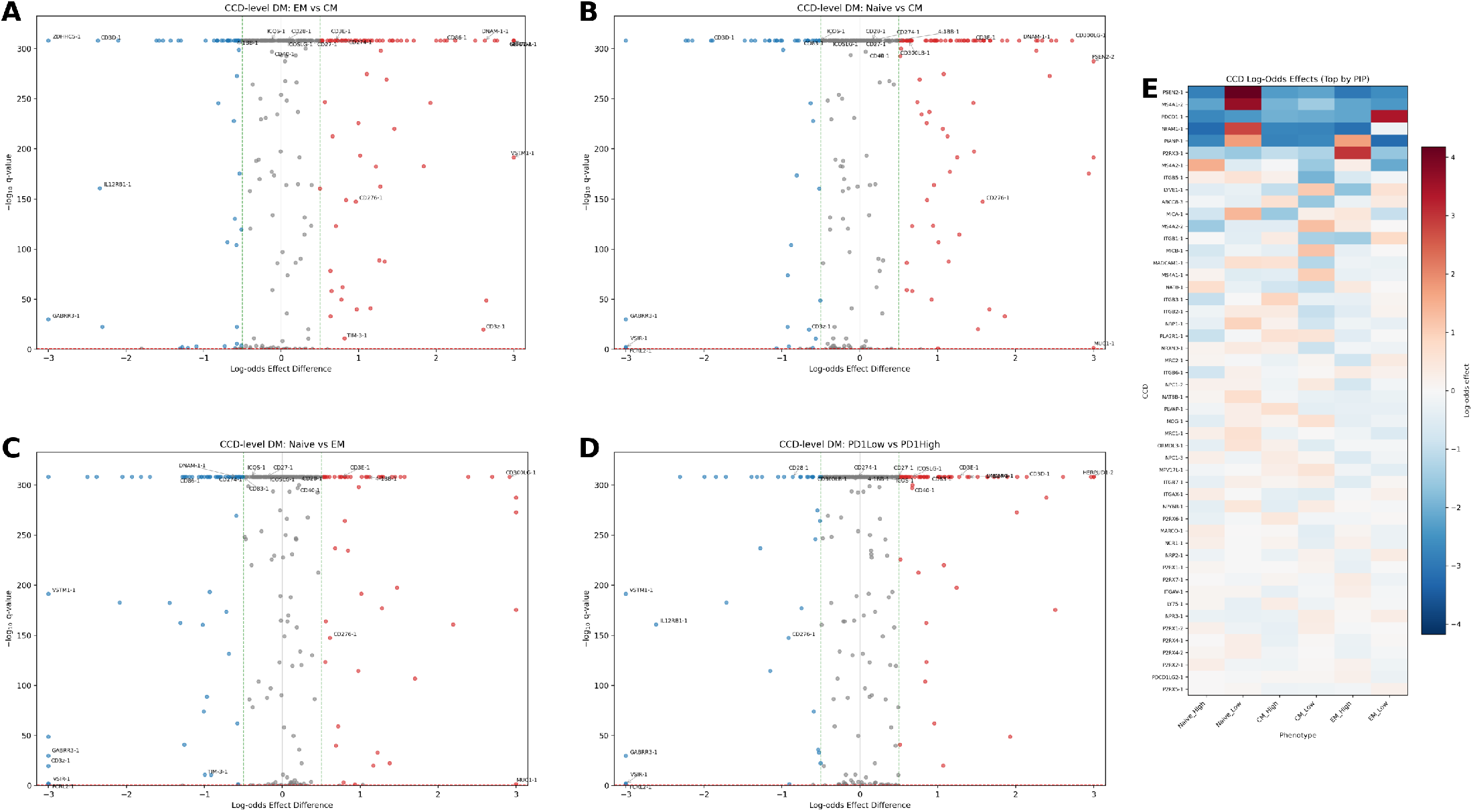
CCD-level analysis reveals combinatorial phenotype effects. (**A–D**) Volcano plots for construct-level phenotype contrasts using leave-one-out G-test. Each point represents one CCD; the x-axis shows log-odds effect relative to the population mean, and the y-axis shows → log_10_(*q*). Points are colored by significance and effect direction at FDR < 0.10. Contrasts shown: (A) EM vs CM, (B) Naïve vs CM, (C) Naïve vs EM, (D) PD-1^low^ vs PD-1^high^. Labels indicate canonical costimulatory domains (4-1BB, CD28, OX40) and other CCDs with large effects. (**E**) Heatmap of log-odds effects across contrasts for CCDs significant in at least one comparison. Unlike ELM-level analysis, numerous CCDs show significant phenotype-shifting effects, demonstrating that specific motif *combinations* determine immunophenotype.

The clinical relevance of these phenotype shifts is direct: CCDs enriched in central memory or naïve compartments would be expected to confer greater persistence and durability, while CCDs enriched in effector memory may enhance immediate cytotoxic function. CCDs associated with low PD-1 expression may reduce exhaustion-related dysfunction. Among canonical costimulatory domains, TNFRSF9 (4-1BB) showed the expected pattern of enrichment in less-differentiated phenotypes (log-odds effect for EM vs CM: →0.93), consistent with its clinical reputation for promoting persistence. CD28 showed the opposite pattern, with enrichment in effector memory (EM vs CM log-odds: +0.34), consistent with its role in driving effector differentiation. ICOS and CD27 also showed significant phenotype-shifting effects, validating the assay against known biology.

### Structural validation: GPCR intracellular domains

To validate that motif-level effects depend on structural context, we analyzed GPCR-derived intracellular domains separately. GPCRs signal primarily through allosteric G-protein activation via structured cytosolic interfaces, with comparatively limited reliance on linear motif-based scaffolding. GPCR intracellular loop 2 (ICL2), which forms a structured G-protein coupling interface, showed minimal CCD-level signal, consistent with its lack of disordered, motif-accessible regions and serving as an internal negative control. GPCR intracellular loop 3 and C-terminal tail (ICL3–CTail), which are more disordered and contain phosphorylation-dependent arrestin recruitment motifs, showed intermediate signal. Non-GPCR domains, which are enriched for intrinsically disordered regions evolved for motif-based scaffold assembly, showed the strongest CCD-level effects.

This pattern supports the interpretation that phenotype-shifting effects require accessible motif combinations capable of engaging downstream signaling scaffolds.

## Discussion

Our central finding is that individual ELMs do not determine CAR T-cell immunophenotype, but specific *combinations* of ELMs do. Mann–Whitney screening identified numerous ELM-phenotype associations, yet Dirichlet-Multinomial modeling, which properly accounts for the compositional structure of FACS-partitioned data, revealed no significant ELM-level effects on phenotype distribution. This discrepancy is informative: single motifs appear to influence proliferation or survival (affecting total construct abundance within phenotypic compartments) without altering the phenotype composition itself. In contrast, CCD-level analysis identified specific costimulatory domains with significant phenotype-shifting effects, demonstrating that it is the particular combination of motifs within a construct that determines differentiation fate.

This finding aligns with fundamental principles of cellular signaling. T-cell differentiation is not determined by any single receptor or motif but emerges from the integration of multiple signaling inputs: strength, duration, and context of TCR signaling; cytokine milieu; metabolic state; and costimulatory signals. ^**? ?**^ It would be surprising if a single ELM could override this integrative logic. Instead, our results suggest that ELMs function as vocabulary elements that, when combined in particular ways, can bias the integrated signaling output toward specific phenotypic outcomes.

### Implications for CAR costimulatory engineering

The practical implication is clear: rational CAR costimulatory engineering should focus on motif combinations rather than individual motifs. The clinical relevance of the identified CCDs is direct. Constructs enriched in central memory or naïve compartments would be expected to confer greater persistence and durability, the key limitation of current CAR T-cell therapies ^4;5^ Constructs enriched in effector memory may enhance immediate cytotoxic function, potentially valuable for aggressive disease. CCDs associated with low PD-1 expression may reduce exhaustion-related dysfunction, though PD-1 is an imperfect proxy for functional exhaustion state.

The CCD-level results provide a starting point for identifying candidate motif combinations. 4-1BB (TNFRSF9) and CD28 showed expected behaviors, validating the assay, while numerous other CCDs exhibited phenotype effects that merit further investigation. Future work could systematically test synthetic constructs combining motifs from CCDs with complementary phenotype profiles (for example, combining motifs associated with CM enrichment and low PD-1 expression).

### Structural context constrains functional accessibility

The stratification of results by GPCR versus non-GPCR origin provides internal validation of the motif-based framework. GPCR intracellular loops 1–2, which form structured G-protein coupling interfaces with minimal accessible linear motifs, showed virtually no CCD-level signal and function as internal negative controls. Non-GPCR domains, evolved for motif-based scaffold assembly at the immune synapse, showed the strongest effects. GPCR ICL3 and C-terminal regions, which are more disordered and contain phosphorylation-dependent arrestin recruitment motifs, showed intermediate signal.

This pattern confirms that phenotype-shifting effects require accessible motif combinations capable of engaging downstream signaling scaffolds, and that the mere presence of sequence motifs is insufficient without the structural context for functional engagement.

### Relationship to prior CAR screening studies

Several recent pooled and combinatorial CAR screening studies have expanded the intracellular signaling design space, but they differ from the present approach in important ways. ^33–36^ Most prior studies use synthetic domains generated by random motif permutation and evaluate aggregate fitness or persistence rather than immunophenotype distributions. By screening evolutionarily selected domains and quantifying effects across defined phenotypic compartments, our approach preserves native motif co-occurrence context and enables statistical inference about which combinations drive differentiation.

The key methodological insight is the distinction between abundance-based and composition-based statistical models. Studies that model total construct abundance (or derived fitness metrics) cannot distinguish proliferation effects from true phenotype-shifting effects. The Dirichlet-Multinomial framework directly addresses this by testing whether a feature alters the *distribution* of cells across phenotypes, independent of how many total cells are present.

### Limitations and future directions

Several limitations warrant consideration. First, the screen measures immunophenotype during ex vivo manufacture, which is an imperfect proxy for in vivo persistence and function. Phenotype associations identified here require validation in models of CAR T-cell persistence and anti-tumor efficacy. Second, our analysis cannot distinguish whether ELMs affect the rate of phenotype acquisition (differentiation kinetics) versus steady-state phenotype maintenance. Third, the current analysis treats motif combinations implicitly (at the CCD level) rather than explicitly modeling motif-motif interactions. Fourth, collapsing ELMs by interacting partner may obscure heterogeneous effects within groups if different binding modes or signaling contexts produce opposing phenotype associations; analysis at the level of individual ELMs will address this limitation. Future work with larger construct libraries could enable formal interaction modeling.

Despite these limitations, the framework provides a principled basis for hypothesis generation. The identified CCDs represent specific, testable motif combinations that can be validated individually and used as starting points for synthetic construct design. A longer-term goal is to develop predictive models that estimate phenotype propensity from ELM composition alone, enabling in silico screening of novel CCDs before experimental validation.

## Conclusion

We screened 1,243 evolutionarily selected intracellular domains in primary human CAR T cells and identified a fundamental distinction: individual ELMs affect CAR T-cell proliferation but do not determine immunophenotype, whereas specific *combinations* of ELMs, represented by individual CCDs, do shift phenotype distributions. This finding suggests that CAR costimulatory engineering should focus on motif combinations rather than single motifs, and provides a statistical framework for identifying candidate combinations from pooled screening data. The identified CCDs represent testable hypotheses for rational design of next-generation costimulatory domains optimized for persistence, effector function, or exhaustion resistance.

## Acknowledgments

This research was supported by the Intramural Research Program of the National Heart, Lung and Blood Institute, National Institutes of Health (NIH). The contributions of the NIH author(s) are considered Works of the United States Government. The findings and conclusions presented in this paper are those of the author(s) and do not necessarily reflect the views of the NIH or the U.S. Department of Health and Human Services.

SC thanks the Aspen Center for Physics for its hospitality and for providing an environment conducive to collaboration. This work was performed in part at the Aspen Center for Physics, which is supported by National Science Foundation grant PHY-2210452.

The anti-CD20 CAR plasmid was a gift from Michael C. Jensen. psPAX2 and pMD2.G were gifts from Didier Trono (Addgene plasmids #12260 and #12259).

Computational work used resources of the NIH HPC Biowulf cluster (https://hpc.nih.gov). We acknowledge the NIH HPC staff for maintaining and supporting the computational infrastructure used in this research.

