## Supplemental File for "A motif-vocabulary model of CAR T-cell intracellular domains identifies determinants of immunophenotype differentiation" (from another version of this preprint: version 1, 2026-02-03)

### Supplementary Methods

**5 Supplementary Figures** **16**

### 1 Data Generation and Experimental Design

This section provides a detailed description of how the screening data were generated, including important considerations about confounding factors and analytical assumptions.

#### 1.1 Lentiviral Library Construction

We synthesized a pooled library of lentiviral transfer plasmids encoding anti-CD20 chimeric antigen receptors (CARs) with variable candidate costimulatory domains (CCDs). All constructs shared identical components except for the CCD:

- Anti-CD20 single-chain variable fragment (scFv) for antigen recognition
- IgG-derived hinge region
- CD28 transmembrane domain
- **Variable CCD** positioned between the transmembrane domain and CD3$\zeta$
- CD3$\zeta$ activation domain
- T2A ribosomal skip sequence followed by truncated EGFR (EGFRt) selection marker

The frequency of each CCD in the lentiviral pool was *not uniform*. As a proxy for CCD frequency in the viral pool, we obtained sequencing data from the transfer plasmid pool. However, this measurement is twice removed from the ideal measurement:

1. **Packaging efficiency**: The efficiency of producing functional lentivirus from different CAR constructs varies. CCD length and sequence can affect lentiviral packaging efficiency.
2. **Transduction efficiency**: The efficiency of transducing T cells with different viral particles varies. CCD identity may affect transduction efficiency through mechanisms including viral titer, receptor binding, and post-entry steps.

#### 1.2 T-Cell Transduction and Enrichment

The pooled lentiviral library was transduced into $CD8^+$ T cells from two healthy donors (constituting two biological replicates). We deliberately targeted a low multiplicity of infection (MOI) of 0.1–0.2 to ensure that the majority of transduced cells received only one CAR construct; transduction of multiple constructs into a single cell would substantially complicate analysis.

Following transduction, $CAR^+$ T cells were enriched by immunomagnetic sorting for cells expressing the truncated EGFR marker. This enrichment step may introduce additional (though likely smaller) perturbations to CCD frequencies.

**Analytical implication**: Given the uncertainty in initial CCD frequencies arising from variable packaging efficiency, transduction efficiency, and enrichment bias, our analysis is limited to questions that are insensitive to the initial distribution of CCDs. The phenotype-stratified design (FACS partitioning within each CCR) specifically enables such analyses.

#### 1.3 Experimental Conditions and Co-Culture Replicates

Cells from each donor were divided into four separate aliquots and subjected to one of four experimental conditions:

1. **Alone**: Cells cultured without stimulation
2. **Activated**: Cells activated nonspecifically with anti-CD3/CD28 antibody-coated beads
3. **K562**: Cells co-cultured with K562 cells (not targets of the anti-CD20 CAR)
4. **Raji**: Cells co-cultured with $CD20^+$ Raji cells (targets of the CAR)

Each of these 8 conditions (2 donors × 4 experimental conditions) was immediately split into 3 technical replicates, yielding 24 *co-culture replicates* (CCRs). This design structure is critical for understanding the correlation structure in the data:

- CCRs from the same donor share transduction history and donor-specific biology
- Technical replicates from the same donor-condition combination share stimulation history
- All samples within a CCR share clonal expansion dynamics during the 10-day co-culture

#### 1.4 FACS Sorting into Immunophenotypic Subfractions

After 10 days of co-culture, each of the 24 CCRs was sorted by fluorescence-activated cell sorting (FACS) into 6 immunophenotypic subfractions (IPSFs) defined by:

- **T-cell subset** (3 levels): Naïve ($CD45RO^-CCR7^+$), Central Memory (CM; $CD45RO^+CCR7^+$), Effector Memory (EM; $CD45RO^+CCR7^-$)
- **PD-1 status** (2 levels): High, Low

This yields 6 IPSFs per CCR: $\text{Naïve}_{\text{High}}$, $\text{Naïve}_{\text{Low}}$, $\text{CM}_{\text{High}}$, $\text{CM}_{\text{Low}}$, $\text{EM}_{\text{High}}$, $\text{EM}_{\text{Low}}$.

**Key analytical insight**: FACS partitions the cells *within each CCR* into the phenotypes of interest. Because this partitioning occurs *after* proliferation and *within* each CCR, phenotype-specific analyses are insensitive to differential proliferation rates across CCDs (which would affect total counts but not within-CCR phenotype distributions).

#### 1.5 Sequencing Library Preparation

From each IPSF, genomic DNA was extracted and used to prepare targeted sequencing libraries. The CCD region was amplified by PCR, followed by addition of Illumina adapter sequences. Approximately uniform amounts of DNA were used from each IPSF.

The final dataset comprised:

$$144 \text{ sequencing libraries} = 2 \text{ donors} \times 4 \text{ conditions} \times 3 \text{ replicates} \times 6 \text{ phenotypes}$$

Although every attempt was made to balance sequencing input across libraries, the libraries were inevitably sequenced to slightly different depths. This variation in library size is accounted for in the statistical models via library size offsets (NB-GLM) or the compositional structure (Dirichlet-Multinomial).

We make the standard assumption of targeted sequencing that library preparation does not materially perturb the distribution of CCDs.

#### 1.6 Candidate Costimulatory Domain (CCD) Background

##### 1.6.1 Functional Properties

Functional CCDs are peptides that lack intrinsic tertiary structure; their signaling function is mediated by the primary amino acid sequence, which serves as a scaffold for binding other components of intracellular signaling networks.

##### 1.6.2 Eukaryotic Linear Motif (ELM) Annotation

The functional elements within each CCD are characterized by Eukaryotic Linear Motifs (ELMs)—short sequence-defined patterns that mediate protein-protein interactions. Each CCD in our library is annotated with the list of ELMs it contains (stored in the `ELMCategory` column of the metadata).

##### 1.6.3 GPCR vs Non-GPCR Classification

CCDs were classified into two broad categories based on their source proteins:

- **Non-GPCR**: Domains from single-pass transmembrane receptors, adaptors, and scaffolds that signal primarily through motif-based scaffold assembly
- **GPCR**: Domains from G-protein coupled receptors (seven-transmembrane proteins)

Within GPCRs, intracellular domains were further subdivided:

- **ICL1–2**: Intracellular loops 1 and 2, which primarily mediate G-protein coupling through structured interfaces
- **ICL3–CTail**: Intracellular loop 3 and C-terminal tail, which are comparatively more disordered and mediate phosphorylation-dependent recruitment of arrestins and modular signaling proteins

GPCR classification was determined by the `NumICD` column in metadata; values > 4 indicate GPCR origin.

#### 1.7 Biological Objectives

The immunophenotype of CAR T-cell infusion products influences clinical outcomes:

- Higher proportions of naïve and central memory cells are associated with better persistence and durability
- Lower PD-1 expression is associated with reduced exhaustion and improved function

Our primary objective was to identify ELM features that affect the distribution of CAR T cells across the six immunophenotypic fractions. To increase statistical power and biological interpretability, we "coarsened" this question into two orthogonal axes:

1. **T-cell subset effects**: Distribution across Naïve vs CM vs EM (pooled over PD-1 status)
2. **PD-1 effects**: Distribution across PD-$1^{\text{High}}$ vs PD-$1^{\text{Low}}$ (pooled over T-cell subset)

#### 1.8 Potential Confounders

##### 1.8.1 Differential Proliferation

Cells with different CCDs (or ELMs) may proliferate or die at different rates within a given CCR. With the current data, we cannot directly distinguish differential proliferation from differential phenotype acquisition.

However, because FACS partitions each CCR *after* proliferation, the within-CCR phenotype distribution is interpretable: it reflects the phenotypic composition of surviving cells with each CCD, regardless of how many total cells survived. This design feature enables analysis of phenotype associations even in the presence of differential proliferation.

##### 1.8.2 Library Size Variation

Sequencing depth varies across libraries. The NB-GLM accounts for this via a log library size offset; the Dirichlet-Multinomial inherently conditions on total counts within each CCD-CCR combination.

### 2 Statistical Model Comparison

We evaluated four statistical approaches for identifying ELM features associated with T-cell phenotype distributions. This section describes each model and presents simulation results comparing their operating characteristics.

#### 2.1 Model Descriptions

##### 2.1.1 Mann–Whitney U Test on Pearson Residuals

**Residual computation**: For each observation (CCD $d$ in library $s$), we compute Pearson residuals:

$$r_{d,s} = \frac{y_{d,s} - \hat{\mu}_{d,s}}{\sqrt{\hat{\mu}_{d,s}}}$$

where $\hat{\mu}_{d,s} = R_d \cdot C_s / T$ is the expected count under independence, with $R_d = \sum_s y_{d,s}$ (row sum), $C_s = \sum_d y_{d,s}$ (column sum), and $T = \sum_{d,s} y_{d,s}$ (grand total).

**Testing**: Within each phenotype $p$, for each ELM group $j$, we partition CCDs into those containing the ELM ($\mathcal{D}_j^+$) and those lacking it ($\mathcal{D}_j^-$). The two-sided Mann–Whitney $U$ test compares the residual distributions:

$$H_0 : F_j^+ = F_j^- \quad \text{vs} \quad H_1 : F_j^+ \neq F_j^-$$

**Effect size**: Cliff's delta quantifies the probability of stochastic dominance:

$$\delta = \frac{1}{n^+ n^-} \sum_{d \in \mathcal{D}_j^+} \sum_{d' \in \mathcal{D}_j^-} \text{sign}(r_d - r_{d'})$$

**Pooled contrasts**: For T-subset comparisons pooled over PD-1, we average Cliff's delta across PD-1 levels and combine $p$-values using Fisher's method.

**Strengths**: Non-parametric; robust to distributional assumptions; high power for detecting any shift; computationally fast.

**Limitations**: Does not model the compositional constraint; effect sizes are not directly interpretable as log-odds; no formal covariance structure across phenotypes.

##### 2.1.2 Per-Phenotype Negative Binomial GLM

**Model**: For each phenotype $p$ separately:

$$y_{d,s} \sim \text{NB}(\mu_{d,s}, \alpha_p), \quad \log \mu_{d,s} = \beta_0 + \sum_j x_{d,j} \beta_j$$

**Inference**: Standard Wald tests on $\beta_j$.

**Strengths**: Model-based with interpretable log-rate effects; accounts for overdispersion.

**Limitations**: Ignores compositional constraint; treats phenotypes as independent; no library size offset (uses per-phenotype data).

##### 2.1.3 Joint Negative Binomial GLM

**Model**: All phenotypes jointly:

$$y_{d,s} \sim \mathrm{NB}(\mu_{d,s}, \alpha)$$
$$\log \mu_{d,s} = \log L_s + \gamma_{\mathrm{CCR}[s]} + \delta_p + \sum_j x_{d,j}\theta_{j,p}$$

**Inference**: To test whether the effect of ELM group $j$ differs between phenotypes $p$ and $q$, we computed contrasts:

$$\Delta_{j,pq} = \theta_{j,p} - \theta_{j,q}$$

Standard errors were obtained from the model's estimated covariance matrix, with cluster-robust variance estimation clustered by CCR to account for residual within-CCR correlation not captured by the fixed effects. The Wald statistic for each contrast is:

$$Z_{j,pq} = \frac{\hat{\theta}_{j,p} - \hat{\theta}_{j,q}}{\mathrm{SE}(\hat{\theta}_{j,p} - \hat{\theta}_{j,q})}$$

which follows an approximate standard normal distribution under the null hypothesis $H_0 : \theta_{j,p} = \theta_{j,q}$.

To increase biological interpretability, we analyzed memory differentiation independently of PD-1 expression by testing pooled contrasts that average motif effects across PD-$1_{\mathrm{high}}$ and PD-$1_{\mathrm{low}}$ within each memory subset. This yielded four primary comparisons: EM vs CM, CM vs Naïve, EM vs Naïve, and PD-$1^{\mathrm{high}}$ vs PD-$1^{\mathrm{low}}$.

**Aggregated Wald statistics**: To summarize results at the level of individual phenotypes, we computed phenotype-specific scores by averaging the signed Wald statistics from all pairwise comparisons involving each phenotype, with signs oriented such that positive values indicate relative enrichment in that phenotype. Formally, let $Z_j^{(p,q)}$ denote the signed Wald statistic for ELM group $j$ comparing phenotypes $p \neq q$. The aggregated score for ELM group $j$ in phenotype $p$ was defined as:

$$\bar{Z}_j^{(p)} = \frac{1}{|K(p)|} \sum_{(p,q)\in K(p)} Z_j^{(p,q)}$$

where $K(p)$ denotes the set of pairwise comparisons involving phenotype $p$.

**Strengths**: Proper covariance structure for contrasts; accounts for CCR effects and library size; single dispersion estimate across phenotypes.

**Limitations**: Does not directly model the compositional constraint (FACS partitioning).

##### 2.1.4 Dirichlet-Multinomial Model

**Model**: For CCD $i$ in CCR $c$:

$$(n_{ic,1}, \ldots, n_{ic,6}) \mid \boldsymbol{\pi}_{ic} \sim \text{Multinomial}(N_{ic}, \boldsymbol{\pi}_{ic})$$
$$\boldsymbol{\pi}_{ic} \sim \text{Dirichlet}(\alpha \cdot \boldsymbol{\mu}_{ic})$$

with

$$\log \frac{\mu_{ic,p}}{\mu_{ic,\text{ref}}} = \beta_{0,p} + \beta_{\text{CCR}[c],p} + \sum_m x_{i,m}\beta_{m,p}$$

**Inference**: Wald tests on contrasts using the full covariance matrix; posterior inclusion probabilities (PIPs) from empirical Bayes spike-and-slab.

**Strengths**: Directly models the compositional constraint; proper uncertainty quantification; PIPs provide calibrated measure of evidence.

**Limitations**: More conservative (fewer discoveries); computationally more intensive; assumes shared concentration parameter $\alpha$.

#### 2.2 Simulation Study Design

To compare the operating characteristics of these four approaches, we conducted simulation studies using data-generating processes calibrated to the observed data.

##### 2.2.1 Simulation Parameters

- **Number of CCDs**: $n = 369$ (matching non-GPCR count)
- **Number of CCRs**: $k = 6$ (matching Raji condition)
- **Number of ELMs**: $m = 49$ (matching normalized ELM groups)
- **ELM prevalence**: Matched to observed frequencies
- **Baseline phenotype proportions**: Matched to observed marginal distribution
- **Concentration parameter**: $\alpha \in \{10, 20, 50\}$ (varying overdispersion)
- **Total counts per CCD-CCR**: Log-normal with mean and CV matched to data
- **CCR effects**: Normal with SD = 0.2 (matched to observed variance)

##### 2.2.2 True Effect Structure

We simulated data under several scenarios:

1. **Global null**: No ELM effects ($\beta_{m,p} = 0$ for all $m, p$)

2. **Sparse effects**: 5 ELMs with true effects (log-odds $\in \{-0.5, -0.25, 0.25, 0.5\}$)

3. **Dense weak effects**: 20 ELMs with small effects (log-odds $\sim N(0, 0.1^2)$)

4. **Matched to data**: Effect structure estimated from the real data

###### 2.2.3 Performance Metrics

For each simulation replicate, we computed:

- **True Positive Rate (TPR)**: Proportion of true non-null ELMs detected at FDR $< 0.10$
- **False Positive Rate (FPR)**: Proportion of true null ELMs falsely detected
- **Precision**: $\mathrm{TP}/(\mathrm{TP}+\mathrm{FP})$
- **Effect size bias**: Mean$(\hat{\beta} - \beta)$ for true non-null ELMs
- **Effect size RMSE**: $\sqrt{\mathrm{Mean}((\hat{\beta} - \beta)^2)}$

#### 2.3 Simulation Results

##### 2.3.1 Summary Statistics

Across all simulation scenarios, we computed aggregate performance metrics for each method (Table 1):

Table 1: **Summary statistics across simulation scenarios.** Power (TPR), false positive rate (FPR), precision, effect size RMSE, and runtime for each statistical method.

| Method | Power | FPR | Precision | RMSE | Runtime (s) |
|---|---|---|---|---|---|
| Mann–Whitney | **0.900** | 0.028 | 0.371 | 0.146 | 0.13 |
| Dirichlet-Multinomial (PIP) | 0.622 | **0.003** | **0.564** | **0.135** | 0.28 |
| Joint NB-GLM | 0.556 | 0.012 | 0.368 | 0.208 | 4.54 |
| Per-Phenotype NB-GLM | 0.389 | 0.006 | 0.357 | 0.210 | 0.09 |

Mann–Whitney achieved the highest power (0.90) but with correspondingly higher FPR (0.028) and lower precision (0.371). The Dirichlet-Multinomial showed the best precision (0.564) and lowest effect size RMSE (0.135) while maintaining excellent FPR control (0.003), though with lower power (0.622). This trade-off motivates using Mann–Whitney for sensitive screening and DM for precise effect estimation. Computational costs varied substantially across methods (Figure 1): Mann–Whitney and Per-Phenotype NB-GLM were fastest ($< 0.15$s), while the Joint NB-GLM was substantially slower ($> 4$s); the Dirichlet-Multinomial achieved good computational efficiency (0.28s) while providing the best precision.

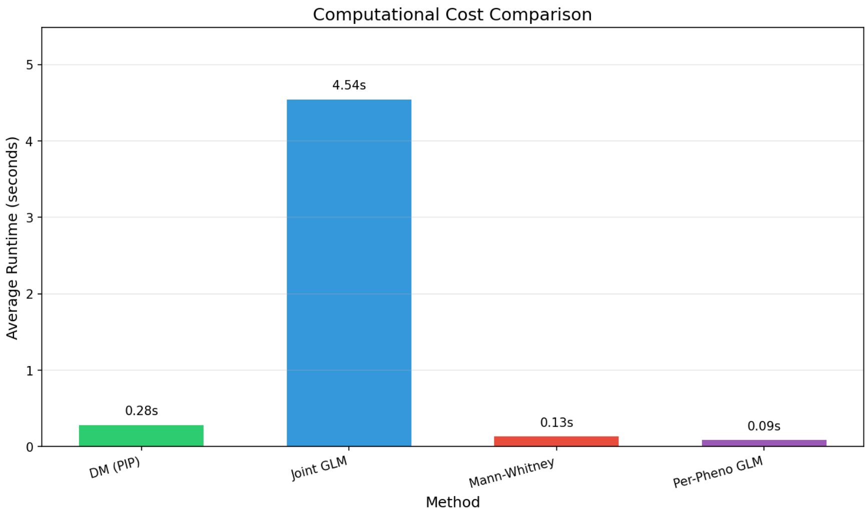

Figure 1: **Computational cost comparison.** Average runtime in seconds for each statistical method. Mann–Whitney and Per-Phenotype NB-GLM are fastest ($< 0.15$s), while Joint NB-GLM is substantially slower ($> 4$s). The Dirichlet-Multinomial achieves good computational efficiency (0.28s) while providing the best precision.

##### 2.3.2 Power by Sample Size and Effect Size

We evaluated statistical power across varying sample sizes ($n = 100, 200, 400$ CCDs) and effect sizes (log-odds $= 0.3, 0.6, 0.9$; Figure 2, Table 2):

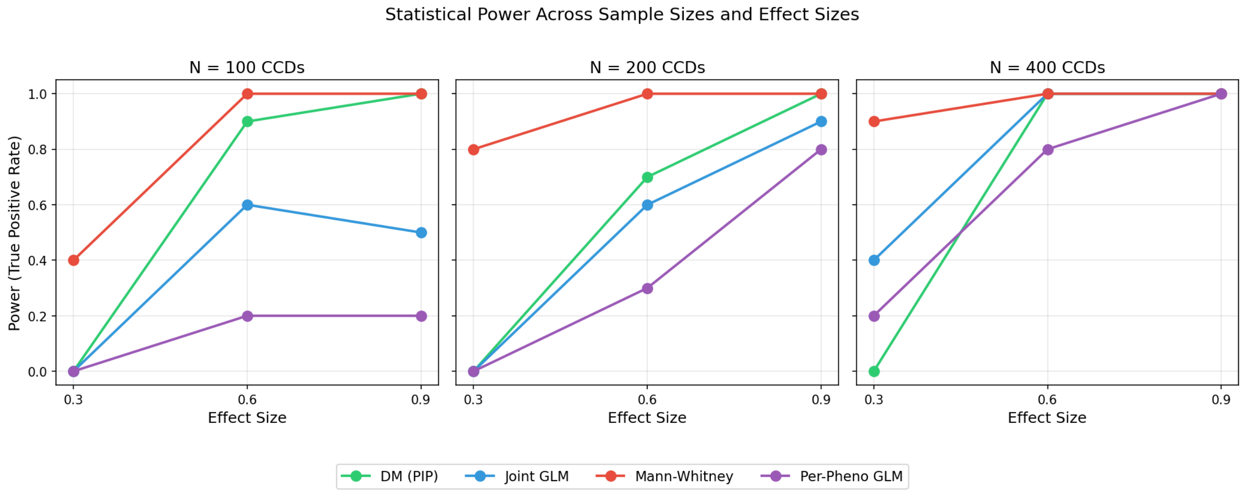

Figure 2: **Statistical power across sample sizes and effect sizes.** Power (true positive rate) as a function of effect size (log-odds) for different sample sizes ($n = 100, 200, 400$ CCDs). Mann–Whitney (red) achieves superior power across all conditions, particularly for small effects. The Dirichlet-Multinomial (green) shows good power for moderate-to-large effects but limited sensitivity to small effects.

Table 2: **Power (TPR) by sample size and effect size.** Each cell shows power for the specified condition.

| | $n$ = 100 | | | $n$ = 200 | | | $n$ = 400 | | |
|---|---|---|---|---|---|---|---|---|---|
| Method | 0.3 | 0.6 | 0.9 | 0.3 | 0.6 | 0.9 | 0.3 | 0.6 | 0.9 |
| Mann–Whitney | 0.4 | **1.0** | **1.0** | **0.8** | **1.0** | **1.0** | **0.9** | **1.0** | **1.0** |
| DM (PIP) | 0.0 | 0.9 | **1.0** | 0.0 | 0.7 | **1.0** | 0.0 | **1.0** | **1.0** |
| Joint NB-GLM | 0.0 | 0.6 | 0.5 | 0.0 | 0.6 | 0.9 | 0.4 | **1.0** | **1.0** |
| Per-Pheno NB-GLM | 0.0 | 0.2 | 0.2 | 0.0 | 0.3 | 0.8 | 0.2 | 0.8 | **1.0** |

Mann–Whitney showed superior power across all conditions, particularly for small effects (0.3) where other methods had near-zero power. At moderate-to-large effect sizes (0.6–0.9), all methods achieved reasonable power with sufficient sample size.

##### 2.3.3 Precision vs Power Trade-off

The precision-power trade-off is visualized in Figure 3.

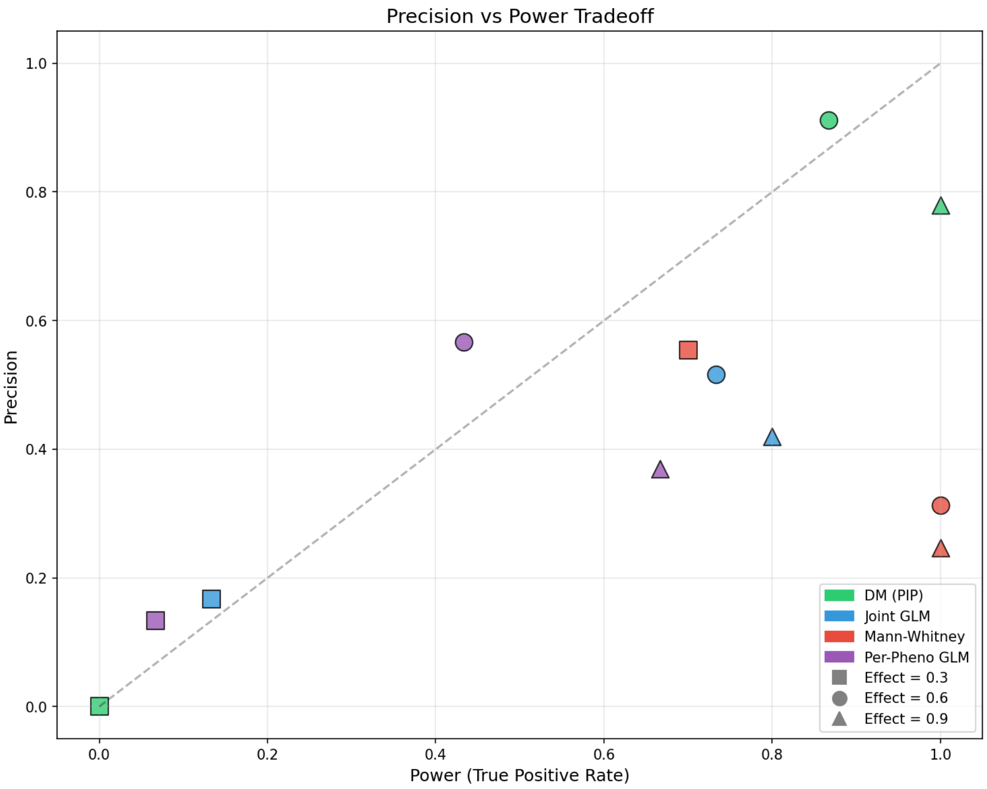

Figure 3: **Precision vs power trade-off.** Precision (y-axis) plotted against power (x-axis) for each method across effect sizes (shape). The dashed diagonal represents ideal balance. The Dirichlet-Multinomial (green) achieves the best precision-power balance, while Mann–Whitney (red) sacrifices precision for maximum power. This trade-off motivates the two-stage approach: MW for screening, DM for refinement.

The Dirichlet-Multinomial achieves the best balance, approaching the diagonal (ideal trade-off), while Mann–Whitney sacrifices precision for power. This motivates the two-stage approach: use Mann–Whitney for sensitive screening, then refine with DM for precise effect estimation.

###### 2.3.4 Effect Size Estimation

For true non-null ELMs, the Dirichlet-Multinomial showed the lowest RMSE (0.135), confirming its suitability for effect size estimation (Figure 4).

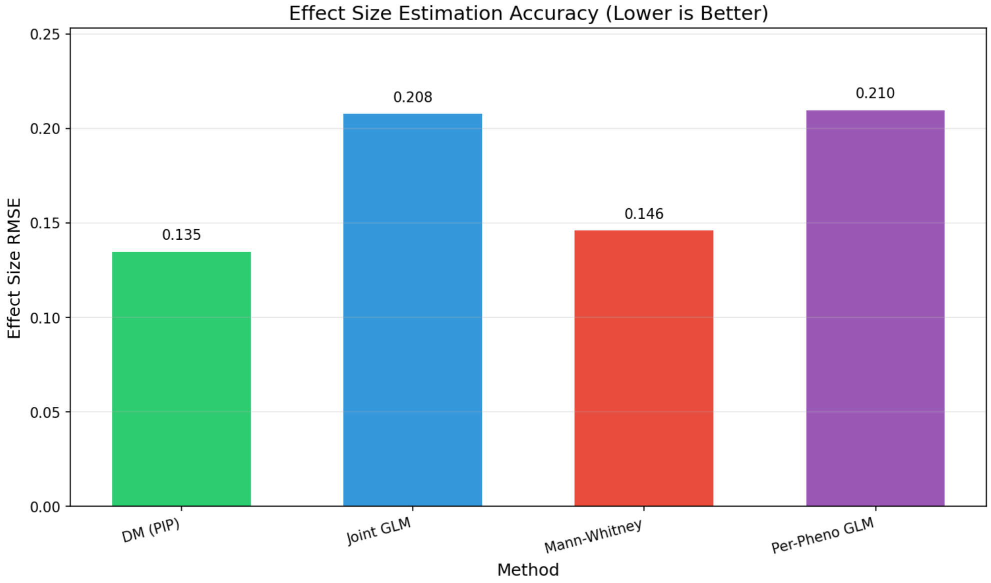

Figure 4: **Effect size estimation accuracy.** Root mean squared error (RMSE) of effect size estimates for each statistical method. Lower values indicate more accurate estimation. The Dirichlet-Multinomial (DM) achieves the lowest RMSE, followed by Mann–Whitney, confirming DM's suitability for effect size refinement.

Note that Mann–Whitney effect sizes (Cliff's delta) are not directly comparable to log-odds effects but showed comparable RMSE (0.146) when appropriately scaled.

##### 2.4 Recommendations

Based on simulation results, we recommend a two-stage approach:

1. **Stage 1: Mann–Whitney screening** at FDR $< 0.10$ for high-sensitivity detection of any ELM-phenotype associations. This stage maximizes power and is appropriate for hypothesis generation.
2. **Stage 2: Dirichlet-Multinomial refinement** for ELMs passing Stage 1. Report PIP $> 0.5$ as high-confidence hits with well-calibrated effect size estimates.

The joint NB-GLM provides a useful intermediate option when formal contrasts are needed but the full compositional structure is not required.

### 3 Sensitivity Analyses

#### 3.1 Robustness to ELM Filtering Threshold

We varied the minimum ELM prevalence threshold from 0.5% to 5% and assessed the impact on:

- Number of ELM groups tested (ranging from 62 at 0.5% to 31 at 5%)
- Number of significant associations at FDR $< 0.10$
- Overlap of top hits across thresholds

The core findings (PDZ, 14-3-3, PP2B, CSL associations) were robust to threshold choice, appearing consistently across all tested values.

#### 3.2 Robustness to CCR Exclusion

We performed leave-one-CCR-out cross-validation to assess the influence of individual CCRs on the results. For each of the 6 CCRs, we re-fit all models excluding that CCR and computed:

- Correlation of effect estimates with full-data estimates
- Number of associations that changed significance status

Effect estimates were highly stable (Pearson $r > 0.95$ for all leave-one-out analyses), and no core findings changed significance status, indicating that results are not driven by outlier CCRs.

#### 3.3 Robustness to Dispersion Estimation

For the NB-GLM and DM models, we varied the approach to dispersion estimation:

- Method-of-moments (default)
- Maximum likelihood
- Fixed values ($\alpha \in \{5, 10, 20, 50, 100\}$)

Results were qualitatively unchanged across estimation approaches, with the core ELM associations remaining significant.

### 4 Supplementary Tables

#### 4.1 Flow Cytometry Antibody Panel

Table 3: **Antibodies used for flow cytometry and cell sorting.** Antibodies marked with † were used for FACS gating and sorting; remaining antibodies were acquired for post-sort characterization.

| Target | Fluorochrome | Manufacturer | Catalog # | Sort† |
|---|---|---|---|---|
| Viability | FVS 440UV | BD Biosciences | 566332 | † |
| CD3 | RB705 | BD Biosciences | 570237 | † |
| CD4 | BV650 | BD Biosciences | 563875 | † |
| CD8 | BV510 | BD Biosciences | 566852 | † |
| IgG (CAR) | Alexa Fluor 647 | Jackson ImmunoResearch | 109-605-088 | † |
| CD45RO | BUV496 | BD Biosciences | 749888 | † |
| CCR7 | APC-Fire 750 | BioLegend | 352926 | † |
| PD-1 | BV711 | BD Biosciences | 564017 | † |
| CD45RA | BUV615 | BD Biosciences | 751555 | |
| CD95 | PE-Cy7 | BioLegend | 305622 | |
| CD28 | Alexa Fluor 700 | BioLegend | 302920 | |
| CD27 | BV605 | BioLegend | 302830 | |
| CD25 | BV785 | BioLegend | 356140 | |
| VISTA | RB744 | BD Biosciences | 757683 | |
| TIGIT | BV480 | BD Biosciences | 747843 | |
| NKG2D | BUV737 | BD Biosciences | 748426 | |

#### 4.2 Full Mann–Whitney Screening Results

Complete results for all 49 ELM groups across all four phenotype comparisons are provided in Supplementary Table S1 (available as `mw_screening_results.xlsx`).

#### 4.3 Dirichlet-Multinomial Refinement Results

Effect estimates, standard errors, $p$-values, and PIPs for all ELMs passing Stage 1 screening are provided in Supplementary Table S2 (available as `dm_refinement_results.xlsx`).

#### 4.4 CCD-Level Results

Individual CCD enrichment scores and PIPs across phenotype comparisons are provided in Supplementary Table S3 (available as `ccd_level_results.xlsx`).

### 5 Supplementary Figures

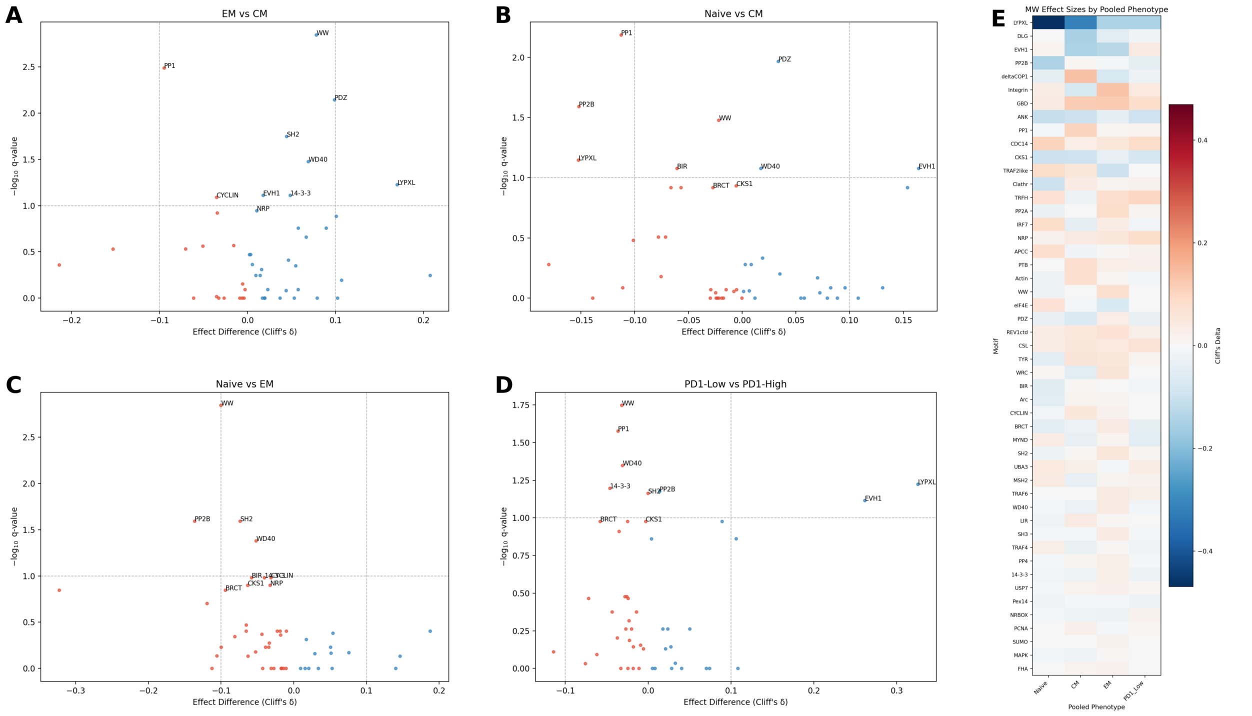

Figure 5: **Mann–Whitney screening identifies ELM-phenotype associations in GPCR ICL3/C-tail domains.** Analysis restricted to ELMs within GPCR intracellular loop 3 and C-terminal tail regions, which are more disordered and contain phosphorylation-dependent arrestin recruitment motifs. (**A–D**) Volcano plots for pooled contrasts comparing ELM effects across immunophenotypes using Mann–Whitney $U$ tests. Each point represents one normalized ELM group. The x-axis shows Cliff's delta ($\delta$) for the contrast $A - B$ (positive values indicate enrichment in phenotype $A$), and the y-axis shows statistical support as $-\log_{10}(q)$, where $q$ is the Benjamini–Hochberg FDR-adjusted $p$-value. Points colored by significance at FDR $< 0.10$. Contrasts shown: (A) EM vs CM, (B) Naïve vs CM, (C) Naïve vs EM, (D) PD-1$^{\text{low}}$ vs PD-1$^{\text{high}}$. (**E**) Heatmap of pooled Cliff's delta values across phenotypes. Compared to the non-GPCR analysis (main text Figure 4), GPCR ICL3/C-tail domains show intermediate signal, reflecting their partial accessibility to linear motif-based signaling.

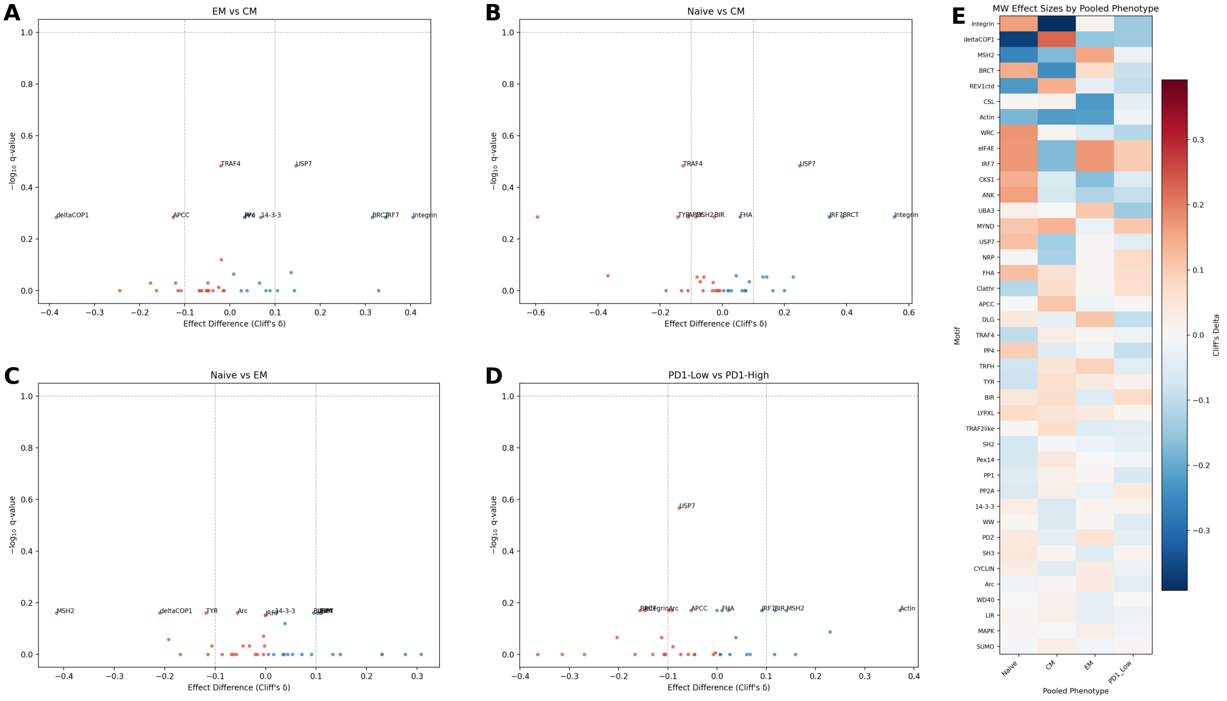

Figure 6: **Mann–Whitney screening in GPCR ICL2 domains (negative control).** (**A–D**) Volcano plots for ELM-phenotype associations in GPCR intracellular loop 2. This structured G-protein coupling interface lacks accessible linear motifs and serves as an internal negative control. As expected, minimal signal is detected, validating that the effects observed in non-GPCR (main text Figure 4) and GPCR ICL3/C-tail domains (Supplementary Figure 1) reflect genuine motif-based signaling rather than technical artifacts. (**E**) Heatmap of pooled Cliff's delta values.

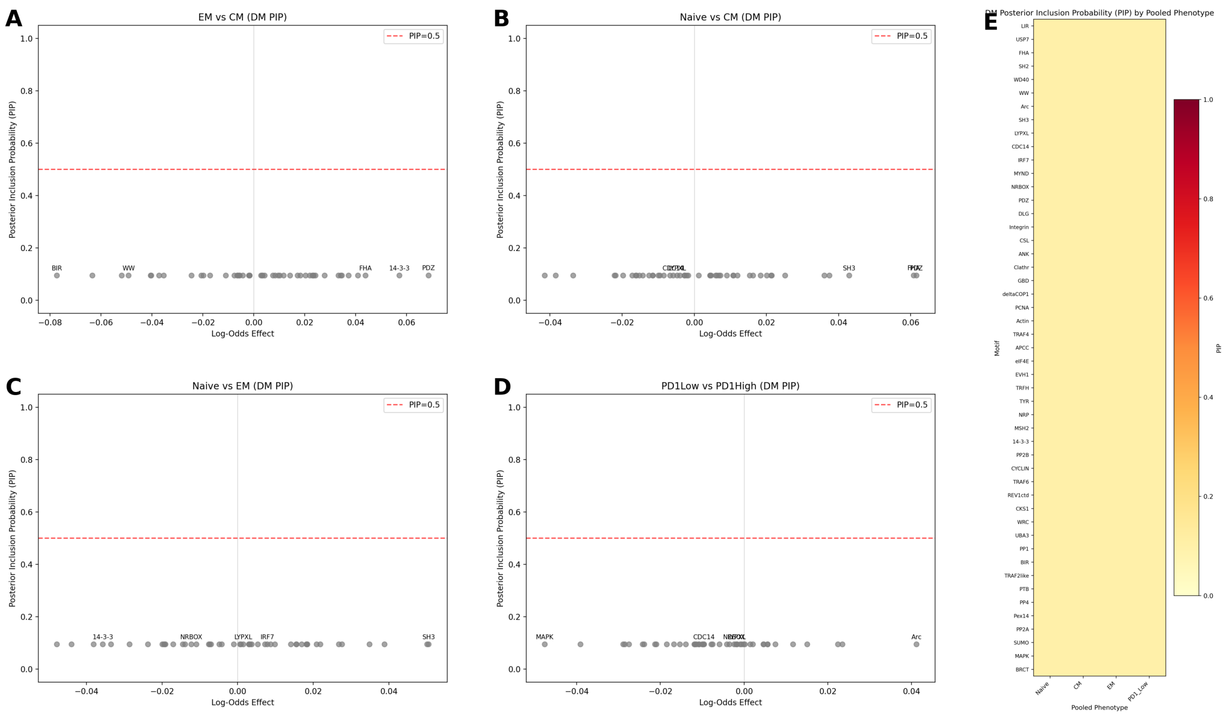

Figure 7: **ELM-level compositional analysis reveals no phenotype effects in non-GPCR CCDs.** Analysis restricted to ELMs within non-GPCR costimulatory domains, which include canonical immune receptors (4-1BB, CD28, OX40, etc.). (**A–D**) Volcano plots for pooled contrasts from the Dirichlet-Multinomial model, which directly accounts for the compositional structure of FACS-partitioned data. The x-axis shows log-odds effect for the contrast $A - B$, and the y-axis shows $-\log_{10}(q)$. Contrasts shown: (A) EM vs CM, (B) Naïve vs CM, (C) Naïve vs EM, (D) PD-$1^{\text{low}}$ vs PD-$1^{\text{high}}$. No ELM-phenotype associations reach FDR significance, indicating that individual motifs do not shift phenotype distributions. (**E**) Heatmap of pooled coefficient Z-scores. This null result, combined with the significant associations detected by Mann–Whitney screening (main text Figure 4), suggests that individual ELMs affect proliferation or survival but not differentiation fate.

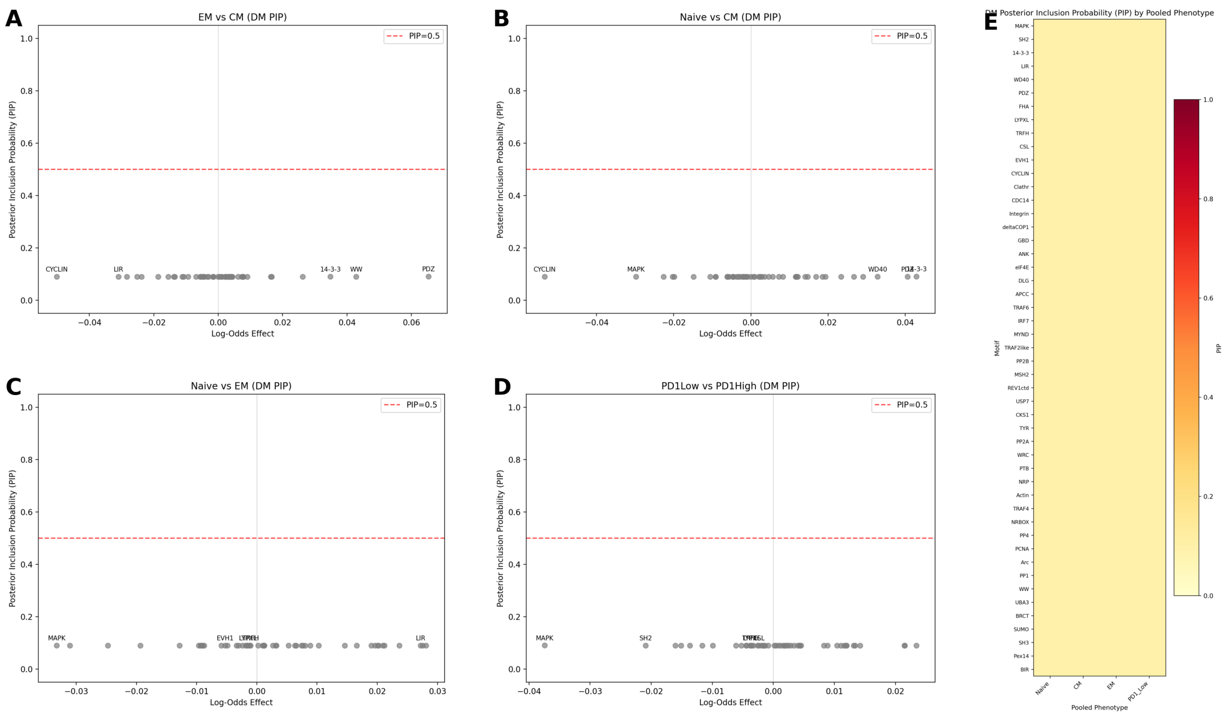

Figure 8: **ELM-level compositional analysis reveals no phenotype effects in GPCR ICL3/C-tail domains.** Analysis restricted to ELMs within GPCR intracellular loop 3 and C-terminal tail regions, which are more disordered and contain phosphorylation-dependent arrestin recruitment motifs. (**A–D**) Volcano plots for pooled contrasts from the Dirichlet-Multinomial model. The x-axis shows log-odds effect for the contrast $A - B$, and the y-axis shows $-\log_{10}(q)$. Contrasts shown: (A) EM vs CM, (B) Naïve vs CM, (C) Naïve vs EM, (D) PD-$1^{\text{low}}$ vs PD-$1^{\text{high}}$. As with non-GPCR domains (Supplementary Figure 3), no ELM-phenotype associations reach FDR significance. (**E**) Heatmap of pooled coefficient Z-scores.

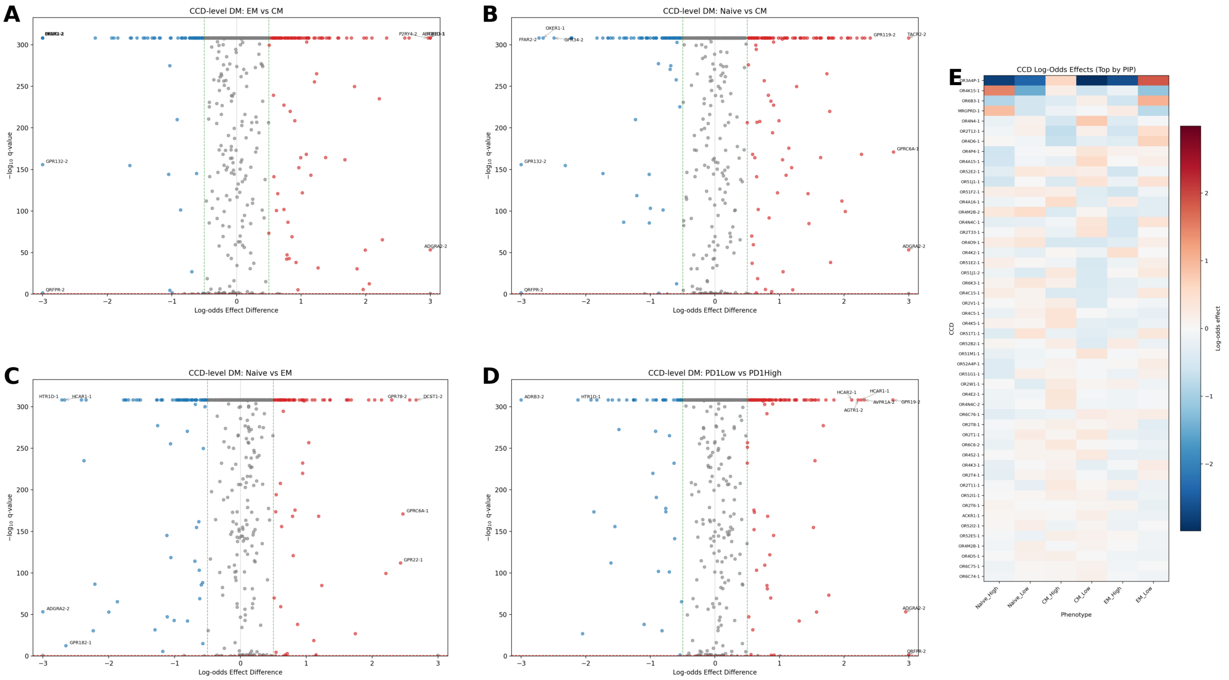

Figure 9: **CCD-level compositional analysis (all GPCR domains).** (**A–D**) Volcano plots for construct-level phenotype contrasts using leave-one-out G-test across all GPCR-derived domains. Each point represents one CCD; the x-axis shows log-odds effect relative to the population mean, and the y-axis shows $-\log_{10}(q)$. Contrasts shown: (A) EM vs CM, (B) Naïve vs CM, (C) Naïve vs EM, (D) PD-$1^{\text{low}}$ vs PD-$1^{\text{high}}$. Intermediate signal compared to non-GPCR domains, reflecting the mixture of structured (ICL2) and disordered (ICL3/C-tail) regions within GPCRs. (**E**) Heatmap of log-odds effects across contrasts.

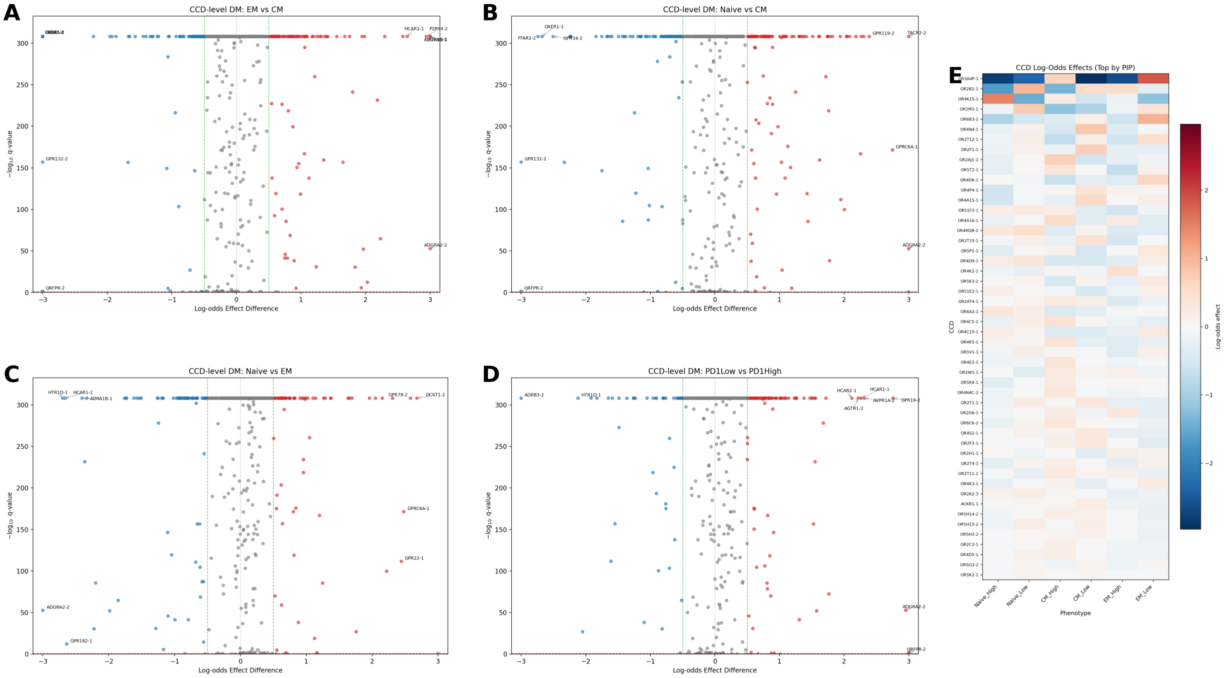

Figure 10: **CCD-level compositional analysis in GPCR ICL3/C-tail domains.** (**A–D**) Volcano plots for construct-level phenotype contrasts in GPCR intracellular loop 3 and C-terminal tail regions using leave-one-out G-test. Each point represents one CCD; the x-axis shows log-odds effect relative to the population mean, and the y-axis shows $-\log_{10}(q)$. Contrasts shown: (A) EM vs CM, (B) Naïve vs CM, (C) Naïve vs EM, (D) PD-1$^{\text{low}}$ vs PD-1$^{\text{high}}$. These disordered GPCR regions show detectable CCD-level phenotype effects, though weaker than non-GPCR domains, supporting the model that accessible motif combinations drive phenotype specification. (**E**) Heatmap of log-odds effects across contrasts.

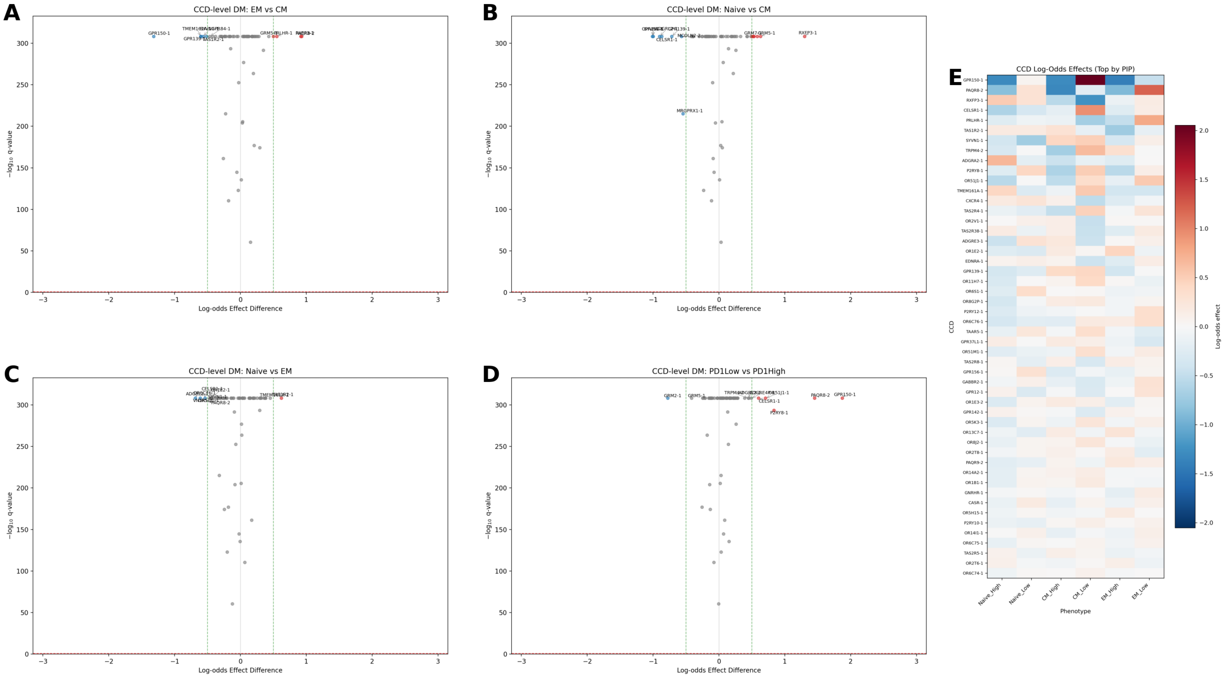

Figure 11: **CCD-level compositional analysis (GPCR ICL2: negative control).** (**A–D**) Volcano plots for construct-level phenotype contrasts in GPCR intracellular loop 2. Each point represents one CCD; the x-axis shows log-odds effect relative to the population mean, and the y-axis shows $-\log_{10}(q)$. Contrasts shown: (A) EM vs CM, (B) Naïve vs CM, (C) Naïve vs EM, (D) PD-1$^{low}$ vs PD-1$^{high}$. The near-absence of significant phenotype-shifting effects in this structured domain provides strong validation that the CCD-level effects observed in non-GPCR domains reflect genuine motif-combination-based signaling rather than technical artifacts. (**E**) Heatmap of log-odds effects across contrasts.
